# Pseudokinase HPO-11 inhibits nonsense-mediated decay to ensure genome stability in *C. elegans*

**DOI:** 10.1101/2022.09.04.506508

**Authors:** Qian Zhao, Erika D Gromoff, Wei Yang, Jennifer Schwarz, Lena Tittel, Ekkehard Schulze, Bettina Warscheid, Ralf Baumeister, Wenjing Qi

## Abstract

DNA double-strand breaks (DSBs) are highly toxic DNA lesions that can induce mutations and chromosome rearrangement therefore causing genome instability (GIN). In response to DSBs, cells activate the DNA damage response by hierarchical assembly of signaling and repair mechanisms. This involves recruitment of the repair factors at DSB sites, local chromatin remodeling, cell cycle arrest and, eventually, DNA repair or apoptosis. Studies investigating mechanisms ensuring genome stability have so far mostly focused on DNA-protein interactions and signal transduction in response to DNA damage. Emerging evidence in the last decade suggests that post-transcriptional control of gene expression by RNA-binding proteins also participates in maintaining genome integrity. However, how specific control of RNA fate mechanistically affects genome stability is still poorly understood. Here, we report that the pseudokinase HPO-11 ensures genome integrity in *C. elegans*. Loss of *hpo-11* leads to accumulation of R-loops, increased DSBs and germline apoptosis, as well as an elevated mutation rate in the somatic cells. In addition, inhibition of nonsense mediated decay (NMD) reduces DSBs and germline apoptosis in the absence of *hpo-11*. We find that HPO-11 physically interacts with SMG-2, the core factor of NMD, and prevents degradation of specific transcripts by NMD, thus contributing to maintenance of genome stability. Furthermore, knock-down of *hpo-11* human homologs NRBP1/2 also results in increased DNA DSBs, and NRBP1/2 physically interact with the human SMG-2 orthologue UPF1. In summary, our work identifies an evolutionarily conserved role of HPO-11 to protect genome stability via suppressing abnormal mRNA decay by NMD.

## Introduction

In eukaryotes, DNA double-stand break (DSB) is the most hazardous DNA damage linked to the occurrence of mutations and chromosomal rearrangement and thus, genome instability (GIN)^1^. GIN severely impairs cellular functions and survival. Proofreading during replication, preventing collision of transcription and replication, as well as DNA damage response (DDR), are critical cellular mechanisms to avoid GIN. Accumulating evidence over the past decade has revealed that defects in DDR contribute to numerous diseases such as premature aging, tumor formation, and neuron degeneration^2^. On the other hand, GIN also plays an essential role in generating genetic diversity. Therefore, a balanced control of GIN must be guaranteed to allow both survival of a species and capability to evolve during natural selection. Previous studies have shown that RNAs and RNA metabolism are both regulators and regulated by DDR to aid in successfully eliminating DNA damages during gene expression^3, 4^. However, the underlying mechanisms, how cells orchestrate the RNA metabolism to protect DNA, are still unclear.

Nonsense mediated decay (NMD) was originally described as an RNA surveillance machinery that is conserved from yeast to human^5^. The core of the NMD machinery composed of several molecules recognizes mRNA transcripts carrying premature termination codons (PTCs) and triggers an RNA degradation pathway to remove these aberrant transcripts. Subsequent research revealed that NMD also plays an important role in post-transcriptional control of eukaryotic gene expression to regulate development, cell cycle progression, and cellular stress responses. In the past, several lines of evidences have suggested that NMD factors are required for the maintenance of genome integrity via affecting multiple aspects of DNA damage response, including DNA replication^6^, telomere maintenance^7^, p53 phosphorylation and G2 checkpoint signaling^8^. In addition, persistent DNA damage may also attenuate specific NMD activities, resulting in stabilization of mRNAs of several critical stress response factors, such as ATF3, to promote survival and adaptation of damaged cells^9^. If stress becomes too severe, NMD may contribute to apoptosis^10^. NMD also negatively regulates homologous recombination (HR) by limiting mRNA and protein levels of key components of HR^11^. All these studies together suggest opposite roles of NMD to affect genome stability that are probably dependent on the cellular context. How these antagonistic contributions of NMD in response to DNA damage are controlled is not yet clear.

Nuclear receptor binding protein 1 and 2 (NRBP1/2) are pseudokinases that are highly conserved across phyla. Recent studies suggest a correlation of expression levels of NRBP1/2 with tumor formation and progression, as well as their modulatory role in regulating diverse signaling pathways, such as AKT, JNK and WNT signaling^12–14^. The only NRBP1/2 homolog in *C. elegans*, HPO-11, contributes to a non-cell-autonomous signaling that promotes tumor formation across tissue boundaries^15^. In addition, HPO-11 was identified as a negative regulator of *lin-12* Notch signaling in regulating vulva development^16^. Mechanistic insights into the apparently diverse functions of NRBP1/2 and HPO-11, however, are mostly missing. To date, the best characterized molecular function of NRBP1 implicates it to be a substrate recognition factor within the Elongin BC E3 ubiquitin ligase complex to regulate proteasome mediated degradation of specific proteins^17, 18^.

In this study we show that HPO-11 prevents GIN in *C. elegans*. Inactivation of *hpo-11* leads to increased accumulation of R-loops and DNA double-stranded breaks (DSBs), resulting in excessive apoptosis in the germline and in an elevated mutation rate in the soma. Germline DSBs and apoptosis, induced by loss of *hpo-11*, is suppressed by the additional inactivation of *smg-2*, encoding a core NMD factor and homolog of UPF1. We found that HPO-11 interacts with SMG-2 to prevent NMD from degrading specific mRNA transcripts that are involved in protection of genome stability. HPO-11, however, does not seem to be a general negative regulator of NMD to degrade transcripts. Moreover, our data suggest that activation of HPO-11 antagonizes SMG-2 in response to genotoxic stress. Furthermore, the human homologs of *hpo-11*, NRBP1 and/or NRBP-2, interact with UPF1. In human cell culture, both NRBP1 and NRBP2 are required to prevent DSBs. In summary, our study suggested a conserved role of HPO-11 and NRBP1/2 in maintaining genome stability probably by preventing degradation of specific mRNAs by NMD.

## Results

### Loss of *hpo-11* results in pleiotropic phenotypes

Our previous study had identified *hpo-11* as a mediator of germline tumor formation that is induced by a non-cell-autonomous function of the FOXO transcription factor DAF-16^15^. *hpo-11* encodes a poorly studied pseudokinase with conserved human homologs that have also been implicated in tumorigenesis. To study functions of *hpo-11*, we generated two *hpo-11* loss of function alleles via CRISPR/Cas9 (Supplementary Fig. 1a). *hpo-11(by227)* was considered as a null allele due to a 6,852 bp deletion that eliminates almost the entire coding region. Consistent with a previous observation^16^, homozygous *hpo-11(by227)* progeny segregated from heterozygous parents were viable, but displayed a pleiotropic phenotype with pale, slowly developed and sterile. The second mutant allele of *hpo-11, by178,* harbors 43 nt insertion at position 46 nt downstream of the start codon, resulting in a PTC in the second exon. *hpo-11(by178)* mutant animals were also pale, showed reduced lipid content, and displayed severe embryonic lethal (Emb) phenotype (Fig. 1a and Supplementary Fig. 1b-1e), suggesting that *hpo-11(by178)* is a hypomorphic mutant. We preferably used *hpo-11(by178)* for subsequent experiments.

**Fig. 1.**
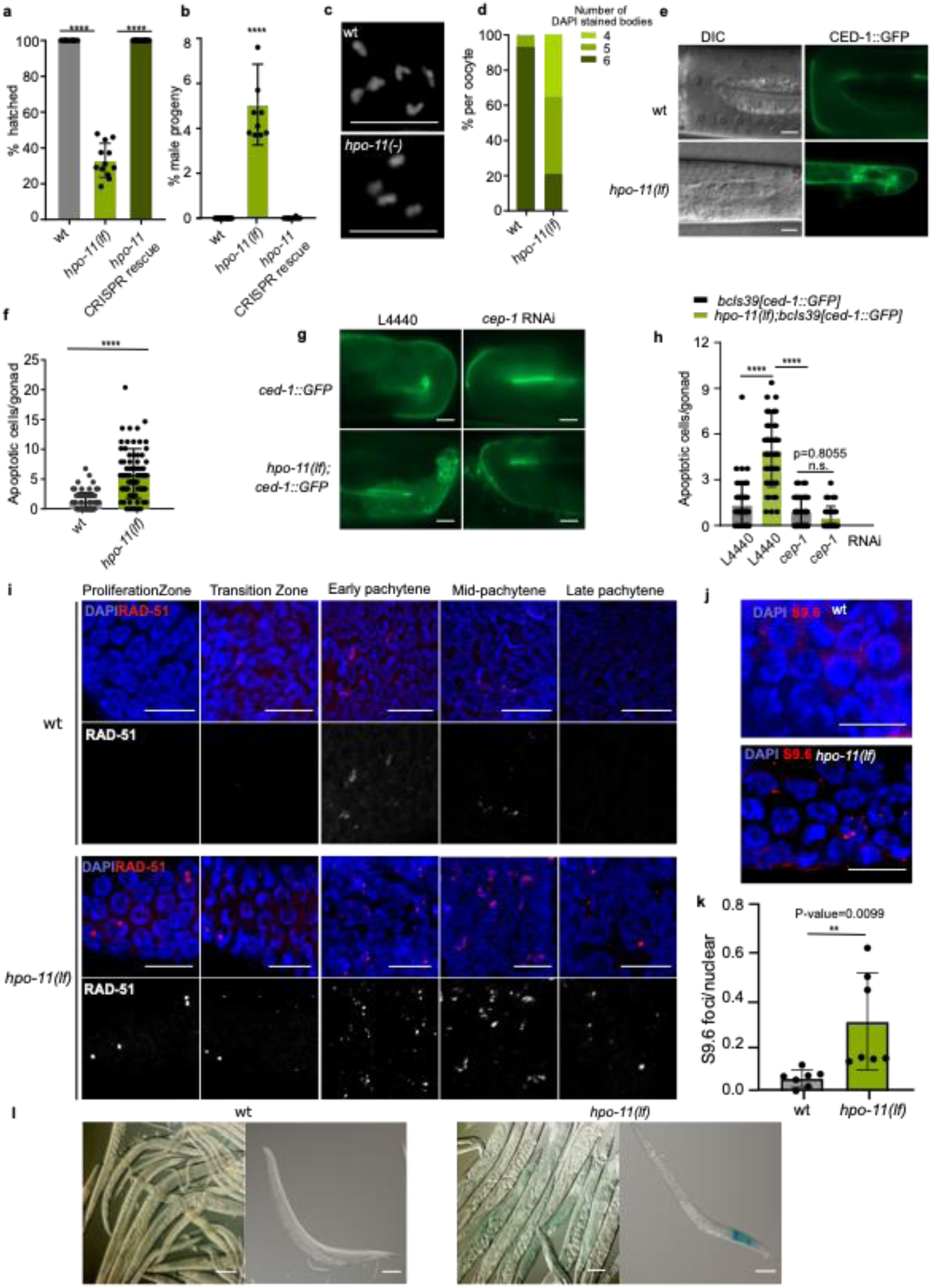
h*p*o*-11* mutation causes genome instability (GIN) a. Embryonic lethal (Emb) phenotype in wild type, *hpo-11(by178)* mutant and *hpo-11(by233)* rescue strains. Percentage of hatched embryos per hermaphrodite: wild type 99.75 ± 0.71 %, n = 10; *hpo-11(by178)* 35.08 ± 9.03 %, n = 12; *hpo-11(by233)* 99.22 ± 1.69, %, n = 16. The data are mean ± SD, statistical test with one-way ANOVA. P values were calculated with Dunnett’s multiple comparisons test. b. Incidence of male progeny in wild type, *hpo-11(by178)* mutant and *hpo-11(by233)* rescue strains. wild type 0.1 ± 0.2 %; *hpo-11(by178)* 5.28 ± 1.94 %; *hpo-11(by233)* 0.1 ± 0.2 %. N = 15 hermaphrodites for each strain, data are mean ± SD, statistical test with one-way ANOVA. P values were calculated with Dunnett’s multiple comparisons test. c. Loss of *hpo-11* leads to reduced number of chromosomes in the oocyte. Shown are representative images of DAPI stained bodies in the most proximal oocytes of wild type and *hpo-11(by178)* mutant animals. Scale bar 5 µm. d. Quantification of the percentage of animals with respective DAPI-staining bodies in the most proximal oocytes for wild type and *hpo-11(by178)* animals. n = 14 for wild type, n = 34 for *hpo-11(by178)*. e. DIC and fluorescence images showing germline apoptotic corpses labeled by CED-1::GFP in wild type and *hpo-11(by178)* mutant animals. Scale bar 10 µm. f. Quantitative analysis of germline apoptotic cells in wild type and *hpo-11(by178)* animals. Number of CED-1::GFP labeled cells in *bcls39[Plim-7::ced-1::GFP]*: 1.11 ± 1.33, n = 78, *hpo-11(by178);bcls39[Plim-7::ced-1::GFP]*: 5.26 ± 3.69, n = 76. Data are mean ± SD, N = 3, statistical test with two-tailed student’s t-test. g. Fluorescence micrographs reveals that activation of *cep-1* contributes to elevated germline apoptosis in *hpo-11(by178)* mutants. Scale bar 10 µm. h. Quantitative analysis of germline apoptotic cells on control RNAi and *cep-1* RNAi. Number of CED-1::GFP labeled cells in wild type + L4440: 1.41 ± 0.22, wild type + *cep-1* RNAi: 0.92 ± 0.15, *hpo-11(by178)* + L4440: 5.29 ± 0.35, *hpo-11(by178) + cep-1* RNAi: 0.51 ± 0.12. Data are mean ± SEM, n > 35, N = 3, statistical test with two-tailed student’s t-test. i. RAD-51 staining of dissected gonads show elevated RAD-51 foci number throughout the gonad in *hpo-11(by178)* worms. At least 5 gonads for each strain. Scale bar 10 µm. j. Representative images show an increased number of S9.6 stained foci in the early pachytene region of the gonad of wild type and *hpo-11(by178)* worms. Scale bar 10 µm. k. Quantification of S9.6 foci that represent R-loops in the early pachytene of wild type, *hpo-11(by178)* day one adult animals. At least five gonads were analyzed. l. *hpo-11(by178);pkIs1604[ATG(A)17GFP::LacZ* mutants show blue patches with X-gal staining. Wild type n = 565, *hpo-11(by178)* n = 459, N = 3 independent biological replicates. Scale bar 100 µm.

### HPO-11 contributes to genome stability

We observed that the few surviving progeny of *hpo-11(by178)* showed a high incidence of male (Him) phenotype (Fig. 1b). Both Him and Emb were recessive and were rescued by eliminating the insertion in the coding region of *by178* via CRISPR/Cas9 that regains a wild type *hpo-11* sequence (allele *hpo-11(by233)*) (Fig. 1a and 1b), indicating that both phenotypic aspects are indeed caused by the insertion in *by178* instead of background mutations. A combination of Emb and Him phenotypes could arise as a consequence of non-disjunction of meiotic chromosomes, derived from failure in meiotic homologous recombination^19^, or may be caused by GIN^20^. To distinguish these two possibilities, we examined chromosome numbers in the oocytes. DAPI staining of the most proximal oocyte of wild type animals shows six paired homologous chromosomes (bivalents)^21^, whereas chromosomal non-disjunction should result in more than six DAPI stained bodies due to presence of unpaired univalent chromosomes. *hpo-11(by178)* animals exhibited either six bivalents or less DAPI stained bodies in the most proximal oocytes (Fig. 1c and 1d), suggesting aneuploidy instead of non-disjunction defects. Next, we asked whether lack of *hpo-11* causes GIN. GIN is typically derived from increased DNA damage that could activate P53 to trigger germline apoptosis and this can be visualized in the strain *bcls39* that expresses a CED-1::GFP reporter to label the initiation of engulfment of apoptotic corpses^22^. *hpo-11(by178)* animals indeed showed significantly more CED-1 labeled cells in the gonad than wild type animals (Fig. 1e and 1f). The GIN induced cell death is distinct from two other types of p53 independent apoptosis in the *C. elegans* germline^23, 24^. One is the physiological apoptosis that eliminates excess oocyte precursors during oogenesis^25^. The other is triggered by pathogen infection and additional environmental stress conditions such as starvation, heat and oxidative stress^23, 24^. We found that RNAi knock-down of p53*/cep-1* almost completely suppressed germline apoptosis in *hpo-11(by178)* animals (Fig. 1g and 1h), suggesting that elevated germline apoptosis in *hpo-11(by178)* might be caused by accumulated DNA damage and GIN. DNA DSB is considered to be the most critical DNA damage to trigger apoptosis. In the *C. elegans* germline, DSB repair is primarily achieved via homologous recombination (HR) that relies on recruitment of the DEAD/H-box RNA helicase RAD-51 to the sites of DSBs. Therefore, appearance of RAD-51-labelled foci serves as a marker for DSBs^26^. In wild type gonad, we observed RAD-51 foci appearing in the transition zone, most frequently in early pachytene, and gradually disappearing in late pachytene, corroborating the previous reports about repair of meiotic crossover derived DSBs via HR^27, 28^ (Fig. 1i and Supplementary Fig. 1f). *hpo-11(by178)* animals not only displayed increased RAD-51 foci in each of these regions, but also showed RAD-51 foci in the proliferation zone, late pachytene and diplotene. Even though we could not distinguish whether the increased RAD-51 foci in the meiotic region were caused by insufficient repair of crossover derived DSBs or, by increased DNA lesion formation, RAD-51 foci in the mitotic proliferation zone are, nevertheless, strong indicators of meiotic crossover independent DSBs. The genome is constantly under threat from endogenous sources such as increased formation of R-loops, a three-stranded nucleic acid transition formed by the nascent RNA, the binding template DNA strand, and the displaced non-template single-stranded DNA during transcription. As DNA transcription and replication share a common template, accumulation of R-loops can result in DNA damage and genome instability when transcription–replication conflicts (TRCs) occur. R-loops also render the non-template single-stranded DNA susceptible to reactive molecules such as reactive oxygen species (ROS)^29^. We wondered whether *hpo-11(by178)* animals have accumulated R-loops. To examine this, we used an antibody S9.6, which recognizes DNA-RNA hybrids, and examined the abundance of positive stained R-loops foci in in the gonad*. hpo-11(by178)* animals had significantly more S9.6 foci than wild type animals (Fig. 1J and 1k), suggesting accumulated R-loops in absence of *hpo-11* that might be the cause of increased DSB formation.

Next, we asked whether loss of *hpo-11* also causes GIN in the soma. In *C. elegans*, DNA DSBs in post-mitotic somatic cells are primarily repaired via non-homologous end joining (NHEJ) which is an error prone repair mechanism^30^. Therefore, increased DSBs formation may result in elevated mutation frequency. To examine the mutation rate in the somatic cells of *C. elegans,* we crossed an out-of-frame *lacZ* reporter *pkIs1604* into *hpo-11(by178)*. Mutations that eventually result in in-frame transformation of the reporter will allow translation of a functional ß-Gal protein and this could be visualized by blue staining in cells in the presence of X-gal^31^. 24% of *hpo-11(lf);pkIs1604* worms showed blue patches, while in *hpo-11(+)* background we barely observed any staining (Fig. 1l), indicating increased mutations rate in the somatic cells that lack of *hpo-11*. Taken together, our data suggest that loss of *hpo-11* causes increased DNA damage, thus resulting in apoptosis in the germline and an elevated mutation rate in the soma.

### Identification of HPO-11-interacting proteins

Like its human orthologues NRBP1 and NRBP2, HPO-11 is proposed to be a pseudokinase due to lack of critical residues in the ATP binding domain^32^ (Supplementary Fig. 2a). As a consequence, these pseudokinases might have modulatory roles as adaptor proteins to mediate protein-protein interactions or as competitors of other kinases for substrate interaction. Identification of HPO-11 protein interactors might therefore provide us valuable hints, how HPO-11 mechanistically contributes to genome stability. For this purpose, we inserted a YPET::LoxP::3xFlag into the genomic *hpo-11* locus using CRISPR/Cas9 genome editing. The resulting knock-in strain *hpo-11(by208[YPET::LoxP::3xFlag::hpo-11])* behaved superficially like wild type animals, suggesting that the fusion protein is functional. Using the anti-Flag antibody we co-immunoprecipitated YPET::LoxP::3xFlag::HPO-11 interacting proteins and analyzed the eluted proteins by quantitative mass spectrometry in a label-free approach. We considered 40 proteins as HPO-11 associated (log_2_ ratio > 2, p < 0.05, n = 3) (Supplementary Data1). Among them, homologs of TSCT-1 (TSC22D3), Y48C3A.12 (TSC22D4) and ELB-1 (Elongin B) have been reported to be interaction partners of NRBP1^17, 33^, confirming the quality of our interactor screen and suggesting a likely functional conservation of HPO-11 and human NRBP1(Fig.2a). Gene ontology (GO) terms analysis using DAVID^34^ revealed that factors regulating development and the reproductive system were enriched among the HPO-11 interactors (Fig. 2b). In addition, the most enriched molecular function of HPO-11 interactor candidates were nucleotide or ribonucleotide binding activity (Fig. 2b), suggesting HPO-11 as a potential regulator of gene expression.

**Fig. 2.**
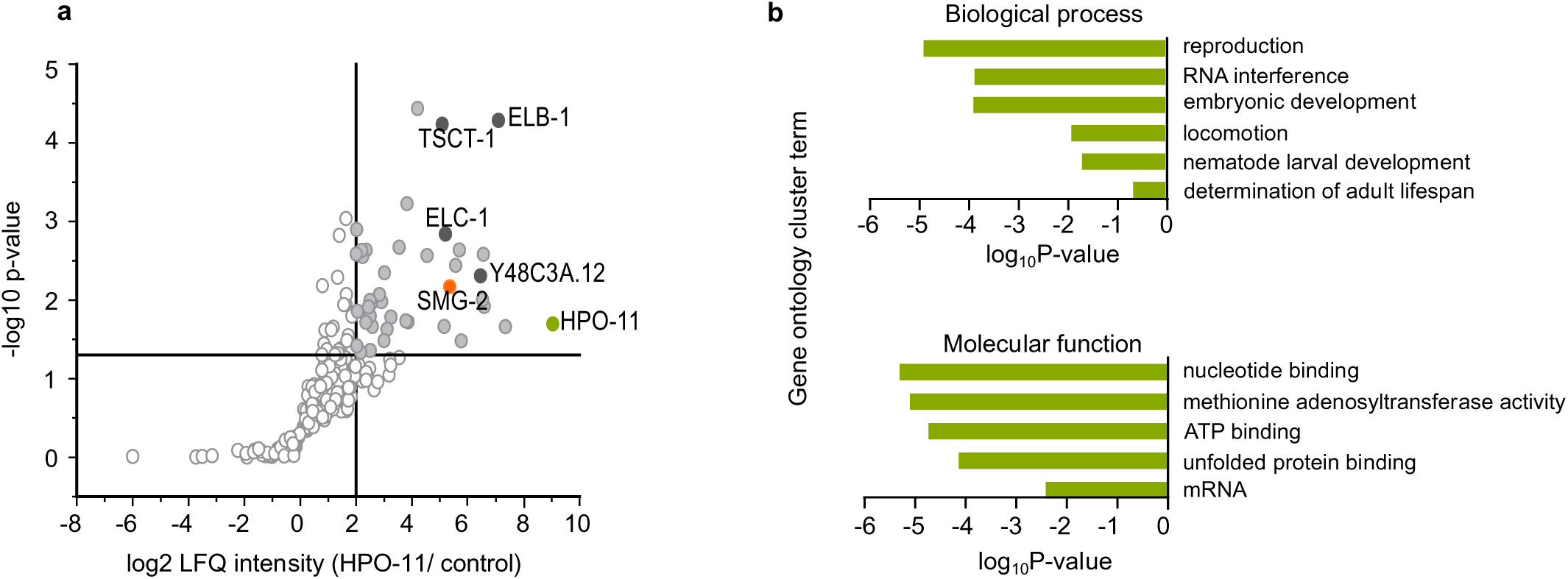
FLAG-HPO-11 interactome a. The difference of log_2_-transformed mean LFQ intensities of FLAG-HPO-11 IP and control IP was plotted against the negative log_10_ p-value of a one-sided two-sample t-test (n=3). Proteins with a p-value below 0.05 and a student’s t-test difference of at least 2 were considered as significantly enriched candidates (light grey). The bait protein HPO-11 is shown in green. Homologs of known NRBP1 interactors (ELB-1, ELC-1 and Y48C3A.12) are depicted in dark grey, SMG-2 in orange. b. Gene ontology term enrichment analysis for the interactors of HPO-11. Selected enriched GO terms are shown with significance.

### HPO-11 ensures genomic stability via inhibiting NMD pathway

Lack of HPO-11 may affect activity of its interactors. Therefore, inactivation of HPO-11 interactors might affect genome stability similarly or oppositely to *hpo-11* inactivation. To identify such factors, we knocked down genes encoding HPO-11 interactors by RNAi and searched for modulation of the *hpo-11(by178)* Emb and Him phenotype. We found that RNAi knockdown of *smg-2* strongly suppressed both Emb and Him phenotypes of *hpo-11(by178)* animals and this was confirmed in *smg-2(qd101) hpo-11(by178)* double mutant animals (Fig. 3a-3d). In addition, *smg-2* RNAi significantly reduced germline apoptosis induced by loss of *hpo-11* (Fig. 3e and 3f). Moreover, *smg-2(qd101) hpo-11(by178)* double mutant animals had less RAD-51 foci from the mitotic to middle pachytene zone than *hpo-11(by178)* mutant animals (Fig. 3g, Supplementary Fig. 1f), even though *smg-2(qd101)* single mutant animals alone displayed more RAD-51 foci than wild type in the meiotic region. This is in agreement with a previous study reporting a possible protective role of NMD against genotoxic stress^35^. Therefore, we conclude that proper adjustments of SMG-2 activity are important to ensure genome stability. Taken together, these observations suggest that HPO-11 may negatively regulate SMG-2 and the excessive SMG-2 activity upon inactivating *hpo-11* causes GIN. Our *hpo-11(by178)* strain used in this study contains a PTC so that *hpo-11(by178)* mRNA might be targeted by NMD for degradation. The 41 nt insertion in *hpo-11(by178)* also results in a frameshift that prevents translation of a functional protein despite of translational read-through^36^. However, in case of an alternative translation start by leaky scanning, stabilization of *hpo-11(by178)* mRNA by *smg-2* mutation might result in translation of a smaller but functional HPO-11 protein that compensate loss of wild type HPO-11 protein^37^. Indeed, *hpo-11(by178)* mRNA level was reduced in comparison to *hpo-11(wt)* mRNA in a *smg-2* dependent manner (Supplementary Fig. S3a), indicating that *hpo-11(by178)* mRNA could be a target of NMD. To test whether *hpo-11(by178)* mRNA could be translated into a functional protein, we generated *hpo-11(wt)::EGFP* and *hpo-11(by178)::EGFP* reporter strains that express the full-length genomic region of either *hpo-11(wt)* or *hpo-11(by178)* fused with EGFP under control of *hpo-11* promoter, respectively. The *hpo-11(wt)::EGFP,* but not the *hpo-11(by178)::EGFP* reporter was actively expressed through out all the development stages (Supplementary Fig. S3b and S3d). *smg-2* RNAi failed to activate expression of *hpo-11(by178)::EGFP*. In contrast, *smg-2* RNAi strongly increased GFP expression level of a PTC-containing GFP::LacZ(PTC) reporter (PTCxi reporter) that does not contain frameshift in GFP codons and thus, could produce a functional GFP protein (Supplementary Fig. S3c). Taken together, these observations suggest that stabilization of *hpo-11(by178)* mRNA upon *smg-2* inactivation does not lead to production of a functional HPO-11 protein. Suppression of the phenotypes in *hpo-11(lf)* animals by inactivating *smg-2* rather indicates a genetic interaction between HPO-11 and SMG-2 to affect GIN.

**Fig. 3.**
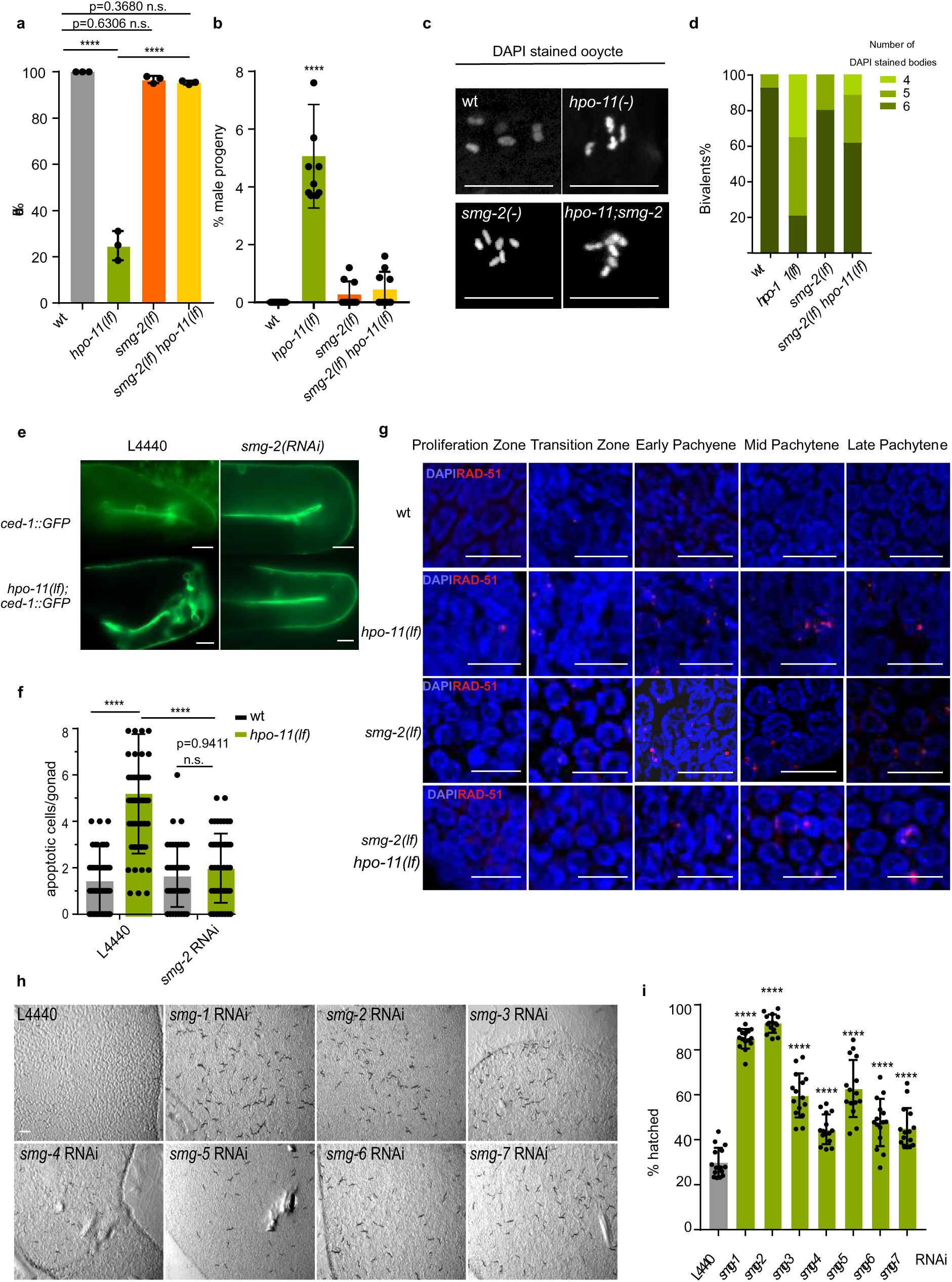
SMG-2 activity in the absence of HPO-11 results in GIN a. Loss of *smg-2* suppresses the Emb phenotype in *hpo-11(by178)* mutants. % of hatched embryos: *hpo-11(by178)* 36.14 ± 9.55%, *smg-2(qd101)* 99.00 ± 0.81%, *smg-2(qd101) hpo-11(by178)* 98.85 ± 1.56. Data are mean ± SD. n = 10 for each strain. Statistical test with one-way ANOVA, P-values were calculated with Tukey’s multiple comparisons test. b. Loss of *smg-2* suppresses the Him phenotype in *hpo-11(by178)* mutants. *hpo-11(by178)*: 5.28 ± 1.94, *smg-2(qd101)*: 0.2125 ± 0.40, *smg-2(qd101) hpo-11(by178)*: 0.44 ± 0.61. Data are mean ± SD. n = 10 for each strain. Statistical test with one-way ANOVA, P-values were calculated with Tukey’s multiple comparisons test. c. Representative images of DAPI stained bodies in the most proximal oocytes in wild type, *hpo-11(by178)*, *smg-2(qd101)* and *smg-2(qd101) hpo-11(by178)* mutant animals. Scale bar 5 µm. d. Quantification of percentage of animals showing respective number of DAPI-staining bodies in the most proximal oocytes. n = 14 for wild type, n = 34 for *hpo-11(by178)*, n = 20 for *smg-2(qd101)* and n = 26 for *smg-2(qd101) hpo-11(by178)*. e. Fluorescence micrographs visualizing germline apoptosis in *hpo-11(by178)* mutants upon *smg-2* RNAi. Scale bar 10 µm. f. *smg-2* RNAi significantly suppresses germline apoptosis in *hpo-11(by178)* animals. Number of CED-1::GFP labeled cells in *bcls39[Plim-7::ced-1::gfp]* + L4440 RNAi control: 1.42 ± 1.57, n = 53; *bcls39[Plim-7::ced-1::gfp]* + *smg-2* RNAi: 1.62 ± 1.31, n = 50; *hpo-11(by178);bcls39[Plim-7::ced-1::gfp]* + L4440 RNAi control: 5.29± 2.57, n = 55; *hpo-11(by178);bcls39[Plim-7::ced-1::gfp]* + *smg-2* RNAi : 1.83 ± 1.70, n = 51; N = 3 biological replicates. Data are mean ± SD, statistical test with One-way ANOVA, P-values were calculated with Tukey’s multiple comparisons test. g. The number of RAD-51 foci in *hpo-11(by178)* gonads are reduced upon *smg-2* RNAi. At least five gonads were analyzed for each strain. Scale bar 10 µm. h. RNAi against *smg-1* to *smg-7* suppress the embryonic lethality of *hpo-11(by178)* to different extents. One *hpo-11(by178)* L4 animals was transferred onto the individual RNAi plates and photographed under transmitted light three days later. Suppression of the Emb phenotype leads to increased number of progenies on the plates. i. Quantification of the percentage of viable embryo of *hpo-11(by178)* feeding with *smg-1* to *smg-7* RNAi bacteria. *hpo-11(by178)* + L4440: 29.81 ± 6.34 %; *hpo-11(by178)* + *smg-1* RNAi: 84.93 ± 4.24 %; *hpo-11(by178)* + *smg-2*RNAi: 91.68 ± 4.10 %; *hpo-11(by178)* + *smg-3* RNAi: 59.73 ± 9.80%; *hpo-11(by178)* + *smg-4* RNAi: 44.64 ± 6.40 %; *hpo-11(by178)* + *smg-5* RNAi: 62.77 ± 12.22%; *hpo-11(by178)* + *smg-6* RNAi: 47.67 ± 10.57 %; *hpo-11(by178)* + *smg-7* RNAi: 45.14 ± 8.97 %. Data are mean ± SD. n = 15 for each RNAi treatment. Statistical test with one-way ANOVA, P-values were calculated with Dunnett’s multiple comparisons test.

*smg-2* encodes a key component of the NMD machinery which is composed of SMG-1-7 proteins. If deletion of HPO-11 activates the NMD to induce GIN, deletion of SMG-1 to SMG-7, like SMG-2, should also result in a suppression of its Emb and Him phenotypes. RNAi inactivation of *smg-1*, *smg-3, smg-4, smg-5, smg-6,* or *smg-7* genes, respectively, suppressed these phenotypic aspects, yet to a variable extent (Fig. 3h and 3i), suggesting that the whole NMD complex contributes to GIN in the absence of *hpo-11*. The variability of suppression may be the consequence of either distinct RNAi knock-down efficacy or a possible functional redundancy of some SMG proteins, which had been proposed by others before^35, 38^.

Our previous genetic screen revealed that inactivation of *hpo-11* suppresses DAF-16 induced germline tumor formation^15^. We asked whether suppression of the germline tumor phenotype upon *hpo-11* knock-down is due to elevated NMD activity. While the germline tumor phenotype of *shc-1;Is[daf-16::GFP]* animals was strongly suppressed in *hpo-11(by178)* mutant background, additional RNAi knockdown of *smg-2* did not result in reoccurrence of germline tumor (Supplementary Fig. 4), suggesting that HPO-11 promotes non-cell autonomous germline tumor formation via an NMD independent mechanism. This result is also an independent confirmation that *smg-2* mutation does not reactivate *hpo-11(by178)* translation via mRNA stabilization.

### HPO-11 interacts with SMG-2 directly and negatively affects NMD activity towards some PTC containing transcripts

NMD is a conserved mRNA surveillance machinery to eliminate aberrant mRNAs which contain PTCs^39^. As our results so far suggest that elevated NMD activity in *hpo-11(by178)* results in GIN, we asked whether HPO-11 negatively regulates NMD activity for degrading PTC-containing transcripts. The PTC-containing GFP::LacZ reporter strain (PTCxi::GFP) has been used to monitor NMD activity^40^. In both wild type and *hpo-11(by178)* background, PTCxi was only weakly expressed, indicating that in both genetic backgrounds NMD is active to degrade this PTC containing reporter. In contrast, *smg-2(qd101)* mutation resulted in strongly enhanced GFP::LacZ expression level, indicating a stabilization of the reporter mRNA (Fig. 4a). Surprisingly, *smg-2(qd101) hpo-11(by178)* double mutant animals also expressed strong but slightly less GFP::LacZ than *smg-2(qd101)* single mutant. Whether this is caused by a NMD independent decay of reporter mRNA or reduced translation in the absence of *hpo-11*, is currently not known.

**Fig.4.**
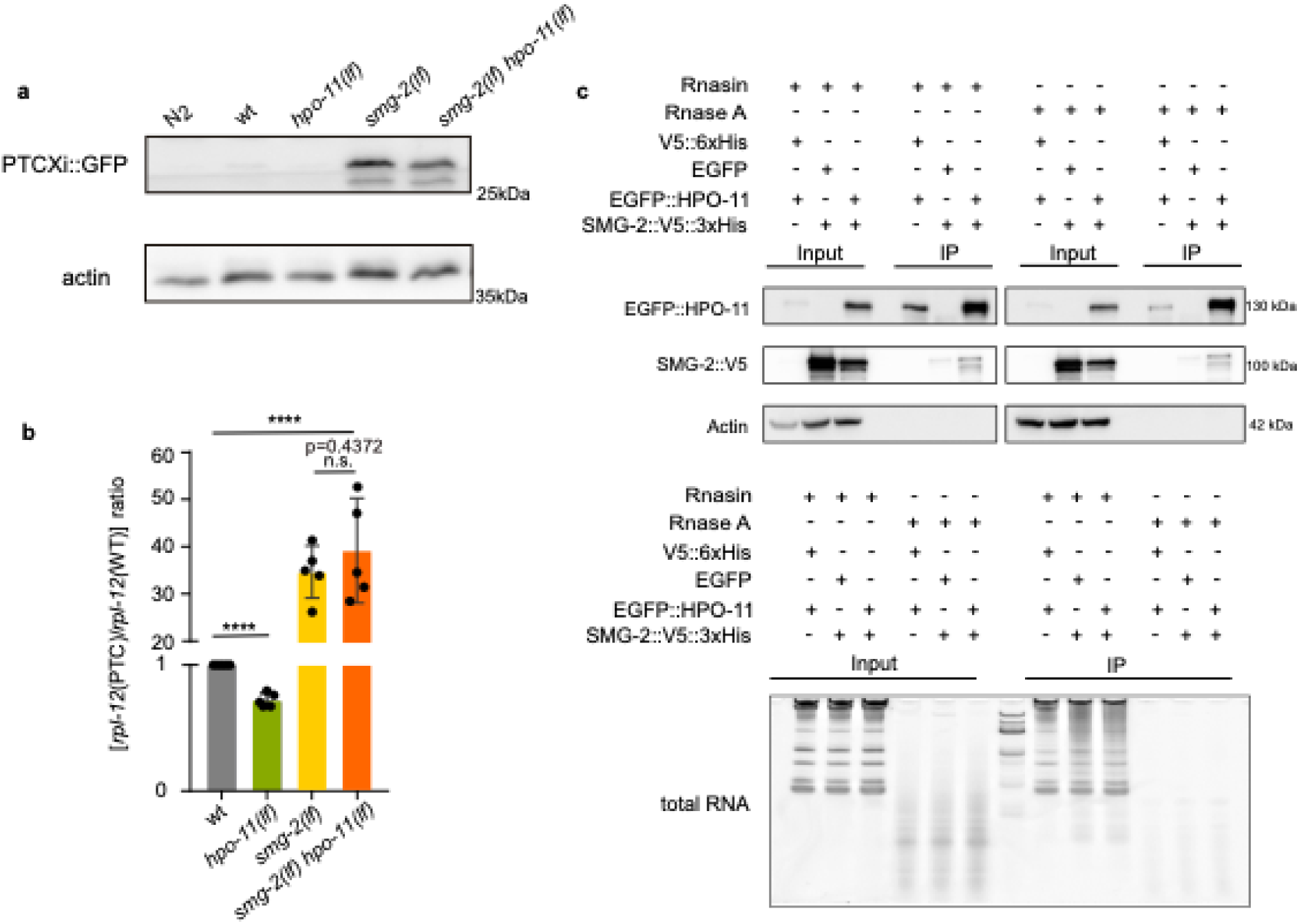
HPO-11 interacts with SMG-2 without affecting general NMD activity a. GFP::LacZ(PTC) protein level in wild type, *hpo-11(by178)*, *smg-2(qd101)* and *smg-2(qd101) hpo-11(by178)* mutant animals. Shown is one representative Western blot detecting GFP::LacZ(PTC). N = 3 biological replicates. N2 (Bristol) without GFP::LacZ(PTC) expression was used as a negative control of GFP::LacZ(PTC) band. b. *hpo-11(lf)* mutant animals have reduced mRNA levels of *rpl-12(PTC) vs. rpl-12(WT)*. Shown are ratios between *rpl-12(PTC)*/*rpl-12(WT)* measured by qRT-PCR. N = 5 independent biological replicates. Statistical tests with two-tailed t test. c. Co-IP of recombinant EGFP::HPO-11 and SMG-2::V5::6xHis proteins in HEK293T cells. Top panel 1-3: Western Blots detecting EGFP::HPO-11, SMG-2::V5::6xHis and actin. Bottom panel: Gel electrophoresis to visualize integrity of the total RNA extracted from the corresponding samples in the presence of RnaseA or Rnase inhibitor (Rnasin). N =3 independent biological replicates for Co-IP. RNAse digestion efficiency was examined in two of three Co-IPs.

Next we tested another NMD target, *rpl-12(PTC)*, which is a product of an alternative splicing event^41^ . RT-qPCR quantification showed a reduction of *rpl-12(PTC)* mRNA in *hpo-11(by178)* (Fig. 4b). In *smg-2(qd101)* animals, *rpl-12(PTC)* mRNA levels displayed an almost 40-fold increase in comparison to wild type. Additional *hpo-11(by178)* mutation did not further alter *rpl-12(PTC)* mRNA level in *smg-2(qd101)* background. Altogether, these results imply that HPO-11 acts as a negative regulator of SMG-2 and, thus, the NMD response for *rpl-12(PTC)*. Finally, we validated the interaction between HPO-11 and SMG-2 observed in the quantitative MS analysis. For this, we expressed EGFP::HPO-11 and SMG-2::V5::6xHis in HEK293T cell line and employed *in vitro* co-immunoprecipitation (coIP) assay to test whether interaction between these two proteins is direct or indirect. With a GFP antibody, we could only co-immuno-precipitate SMG-2::V5::6xHis in the presence of EGFP::HPO-11 but not EGFP alone (Fig. 4c). In addition, RNase treatment led to RNA decay without disrupting interaction between HPO-11 and SMG-2, suggesting a possible RNA independent direct interaction between HPO-11 and SMG-2 (Fig. 4c).

### HPO-11 protects specific transcripts against NMD to ensure genome stability

Our results so far indicate an inhibitory role of HPO-11 on NMD to prevent GIN. To investigate transcripts governed by HPO-11 and SMG-2, we sequenced polyadenylated RNA from wild type, *hpo-11(by178)*, *smg-2(qd101)* and *smg-2(qd101) hpo-11(by178)* double mutant with three independent biological replicates. We obtained an average of 36.3 million reads per sample with a more than 83% aligned rate to the *C. elegans* reference genome (WBcel235.96, Supplementary Fig. 5a). Next, we used DESeq2 package to assess genes with a statistically significant response to *hpo-11*. As the major function of NMD is to degrade mRNAs and our data suggest a hyperactivation of NMD in *hpo-11(by178)* mutant animals, we only focused in this study on genes that were significantly lower expressed in *hpo-11(by178)* than in wild type background. We found that 78 genes were 2-fold down-regulated in *hpo-11(by178) vs.* wild type animals (FC > 2, FDR < 0.05) (Fig. 5a, Supplementary Data 2). In addition, down-regulation of 32 out of these 78 genes upon *hpo-11* inactivation was *smg-2* dependent, suggesting NMD to be responsible for down-regulation of these transcripts in *hpo-11(by178)* mutant background. Additional RT-qPCR of the selected mRNAs confirmed the regulatory antagonism of HPO-11 and SMG-2 (Fig. 5b).

**Fig. 5.**
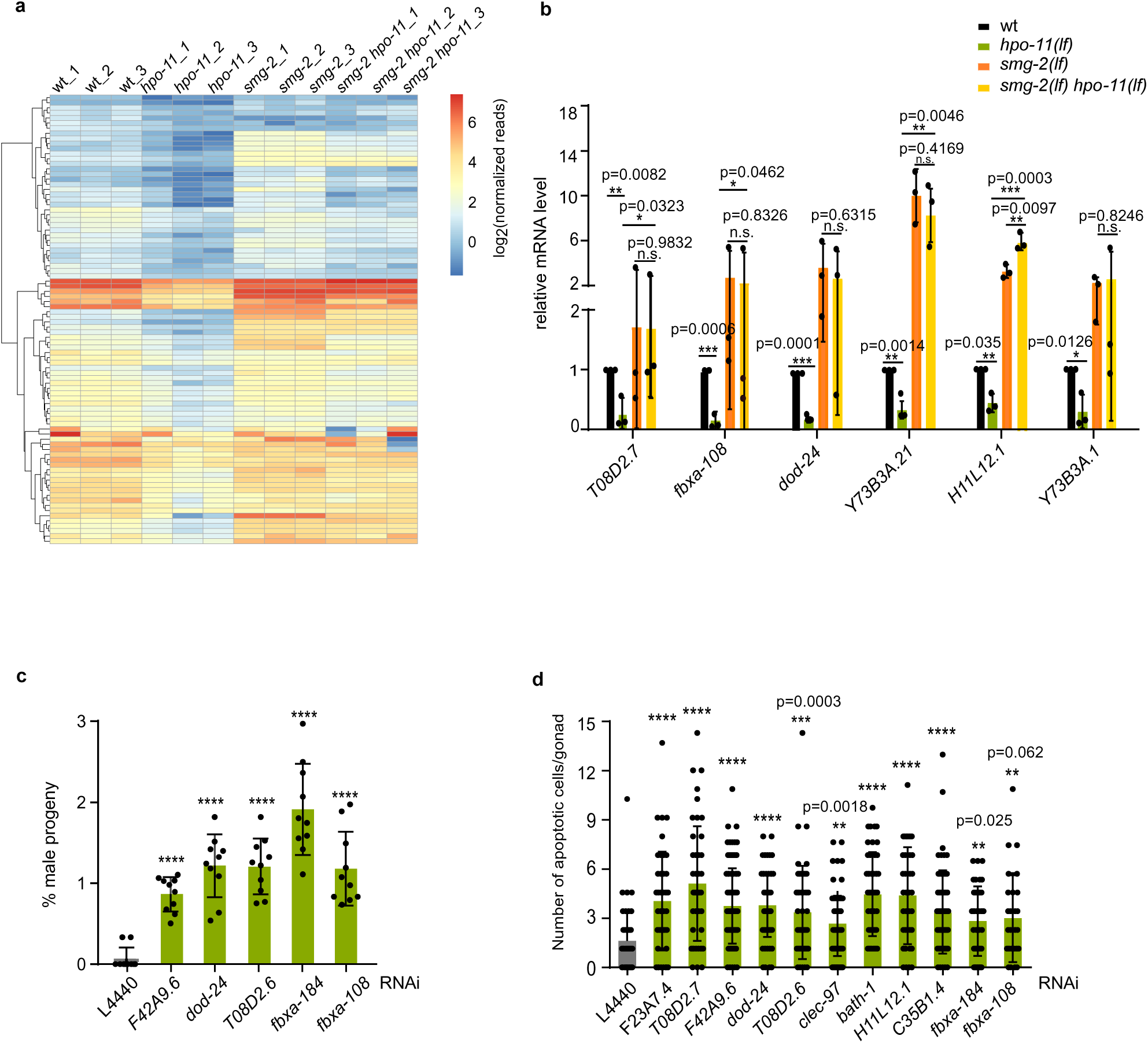
HPO-11 protects specific mRNAs from degradation by NMD machinery a. Heatmap of the 78 genes that are significantly down-regulated (FC > 2 and FDR < 0.05) in *hpo-11(by178) vs.* wild type. b. qRT-PCR of selected mRNA targets that are down-regulated in *hpo-11(by178)* in a SMG-2 dependent manner. N= 3 independent biological replicates. Statistical test with two-tailed student’s t-test. c. RNAi knockdown of 5 NMD targets controlled by HPO-11 results in Him phenotype. L4440: 0.08 ± 0.15 %; *F42A9.6*: 0.93 ± 0.17 %; *dod-24*: 1.17 ± 0.42 %; *T08D2.6*: 1.19 ± 0.39 %; *fbxa-184*: 1.82 ± 0.59 %; *fbxa-108*: 1.18 ± 0.46 %. Data are mean ± SD. n = 10 for each RNAi knockdown. Statistical test with two-tailed student’s t-test. d. RNAi knockdown of 11 NMD targets controlled by HPO-11 causes elevated numbers of apoptotic cell corpses in the germline. L4440 RNAi control: 1.36 ± 1.55, n = 53; *F23A7.4* RNAi: 3.55 ± 2.58, n = 51; *T08D2.7* RNAi: 4.48 ± 3.06, n = 50; *F42A9.6* RNAi: 3.29 ± 2.02, n = 94; *dod-24* RNAi: 3.23 ± 1.70, n = 65; *T08D2.6* RNAi: 2.94 ± 2.49, n = 51; *clec-97* RNAi: 2.34 ± 1.74, n = 91; *bath-1* RNAi: 3.89 ± 2.22, n = 57; *H11L12.1* RNAi: 3.84 ± 2.60, n = 44; *C35B1.4* RNAi: 2.94 ± 2.19, n = 78; *fbxa-184* RNAi: 2.48 ± 1.86, n = 48; *fbxa-108* RNAi: 2.63 ± 2.36, n = 30. N = 3 independent biological replicates. Data are mean ± SD, Statistical test with two-tailed student’s t-test.

Our transcriptome analysis additionally revealed up-regulation of 1816 transcripts in *smg-2(qd101) vs.* wild type day one adult animals (FC > 2, FDR < 0.01, Supplementary Data 2). Compared with a previous RIP-seq analysis identifying mRNAs bound by SMG-2^42^, we found 23% (419/1816) of up-regulated transcripts were SMG-2 bound, suggesting them as possible direct mRNA targets of NMD. This result is in line with the conclusion of the same study^42^ that about 25% of the SMG-2 bound transcripts are regulated by SMG-2. The fact that only 7 of these 419 NMD direct target mRNAs were down-regulated in *hpo-11(by178)* mutants suggests that HPO-11 is not a general negative regulator of the NMD to control gene expression.

Among the genes down-regulated in *hpo-11(by178)* animals by NMD, 22 of the 32 transcripts are germline enriched (Supplementary Fig. 5c). Moreover, *T08D2.7* is a homolog of check point kinase 2 and *fbxa-108* encodes a F-box domain containing protein. Both *T08D2.7* and *fbxa-108* have been implicated in regulation of meiotic chromosomal segregation and chromosomal morphology in the germline^43, 44^. To examine whether down-regulation of these targets alone is sufficient to generate a GIN phenotype, we performed RNAi knock-down of 15 of the 32 genes, for which *E.coli* RNAi strains were available, and examined Him phenotype and germline apoptosis. Knock-down of five out of 15 genes resulted in a mild Him phenotype and RNAi against 11 genes caused increased the number of apoptotic cell corpses in the germline (Fig. 5c and 5d). These data indicate that degradation of these targets by NMD might count for GIN upon inactivation of *hpo-11*.

### HPO-11 regulates response to DNA DSBs

Our results so far suggest a protective role of HPO-11 from endogenous GIN. We further asked whether HPO-11 activity is also involved in response to exogenous sources of DNA damage. The drug Camptothecin (CPT) causes DNA DSBs via preventing relegation of DNA breakages introduced by DNA topoisomerase. CPT treatment causes Emb phenotype in wild type animals in a concentration dependent manner. As *hpo-11(by178)* animals already showed a highly penetrant Emb phenotype without CTP treatment, we applied CPT and reduced *hpo-11* activity by RNA interference that did not cause an Emb phenotype. Exposing wild type L4 animals to 5 µM CPT for 24 h resulted in about 20% Emb in the progeny produced by the day one adult animals (Fig. 6a). *hpo-11* RNAi strongly increased the Emb phenotype after CPT treatment, supporting a protective role of HPO-11 in response to DSBs formation. *smg-2* mutants are RNAi resistant^45^, which did not allow us to test *hpo-11* RNAi knock-down in *smg-2(qd101)* background. Nevertheless, *smg-2(qd101)* animals were slightly more resistant to CPT toxicity and the *smg-2(qd101) hpo-11(by178)* double mutants were significantly more resistant as wild type (Fig. 6b), indicating that NMD activity has a detrimental effect in response to DNA damage.

**Fig. 6.**
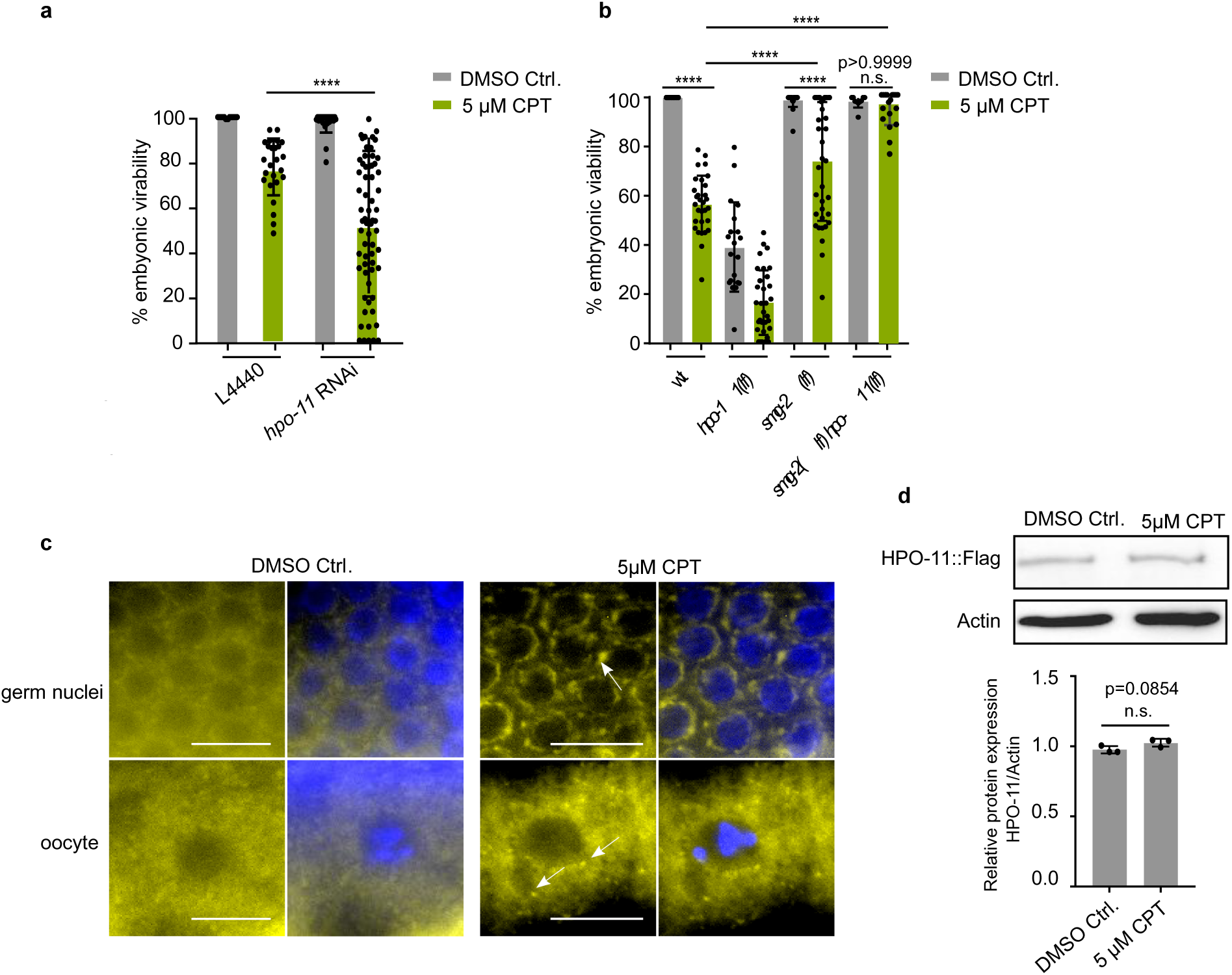
HPO-11 regulates response to CPT induced DNA damage a. RNAi knock-down of *hpo-11* reduces embryonic viability upon 5 µM CPT treatment. Hatched embryo in wild type + L4440 DMSO control: 99.91± 0.22 %, n = 20; wild type + L4440 CPT treated: 75.97 ± 9.80 %, n = 25; wild type + *hpo-11* RNAi DMSO control: 98.31 ± 3.63 %, n = 25; wild type + *hpo-11* RNAi CPT: 48.90 ± 26.78 %. n = 66. Shown are pooled data of 3 biological replicates. Data are mean ± SD. Statistical test with One-way ANOVA. b. CPT hypersensitivity of *hpo-11(by178)* mutant animals is suppressed by additional inactivation of *smg-2*. Hatched embryo in wild type DMSO control: 99.9 ± 0.29 %, n = 11; wild type CPT treated: 56.38 ± 11.66 %, n = 30; *hpo-11(by178)* DMSO control: 38.85 ± 18.30 %, n = 20; *hpo-11(by178)* CPT treated: 16.09 ± 13.22 %, n = 32; *smg-2(qd101)* DMSO control: 99.09± 2.88 %, n = 25; *smg-2(qd101)* CPT treated: 73.88 ± 24.34 %, n = 37; *smg-2(qd101) hpo-11(by178)* DMSO control: 98.35 ± 2.42 %, n = 10; *smg-2(qd101) hpo-11(by178)* CPT treated: 95.33 ± 6.56 %, n = 20. Shown are pooled data of 3 biological replicates. Data are mean ±SD. Statistical test with two tailed Student’s t-test. c. YPET::loxP::3xFlag::HPO-11 protein accumulates at the perinuclear region of germs and oocytes upon CPT treatment. Scale bar 10 µm. d. CPT treatment does not affect the protein level of YPET::loxP::3xFlag::HPO-11. N = 3 replicates. Statistical test with two tailed Student’s t-test.

CPT treatment also seems to result in the spatial redistribution of HPO-11 protein. YPET::LoxP::3xFlag::HPO-11 in our CRISPR-knock-in strain showed broad expression both in the soma and the germline (Supplementary Fig. 6a-e). In addition, YPET::LoxP::3xFlag::HPO-11 was primarily diffusely distributed in the somatic cytosol, but formed punctate structure in the gonad core, around the germ nuclei, as well as in the oocytes. Such punctate structures are probably ribonucleoprotein particles that play an important role in regulating the RNA life cycle. Upon CPT treatment, we observed an increased YPET::LoxP::3xFlag::HPO-11 localization to the perinuclear punctate structures without obvious alterations in protein expression levels (Fig. 6c and 6d). We propose that the physical redistribution of HPO-11 upon DSBs formation might contribute to its protective role against GIN, probably via inhibition of NMD.

### Human NRBP1 and NRBP2 interact with UPF1 and ensure genome stability

NRBP1 and NRBP2 are the two human orthologues of *hpo-11* (Supplementary Fig. 2a) that have been implicated to affect cancer development. To assess a possible functional conservation between NRBP1/2 and HPO-11, we first tested whether NRBP1/2 also interact with UPF1, the mammalian orthologue of SMG-2. We found that endogenous UPF1 co-immunoprecipitated with an antibody recognizing both NRBP1 and NRBP2 proteins in MCF-7 cells (Fig. 7a). Next, we asked whether knock-down of NRBP1 or NRBP2 induces DSBs. Following DSBs formation, histone H2AX becomes phosphorylated, which then is referred to as γH2AX and used as an indicator for DNA DSBs^46^. We found that siRNA knock-down of either NRBP1 or NRBP2 led to an increase in γH2AX level compared to the control, and double knock-down of both NRBP1 and NRBP2 resulted in a further increase of γH2AX, indicating redundant functions of NRBP1 and NRBP2 in preventing formation of DSBs (Fig. 7b). Taken together, these results suggest that both the interactions of NRBP1/2 and HPO-11 with UPF1/SMG-2 and their roles in the control of genome stability are conserved in evolution.

**Fig. 7.**
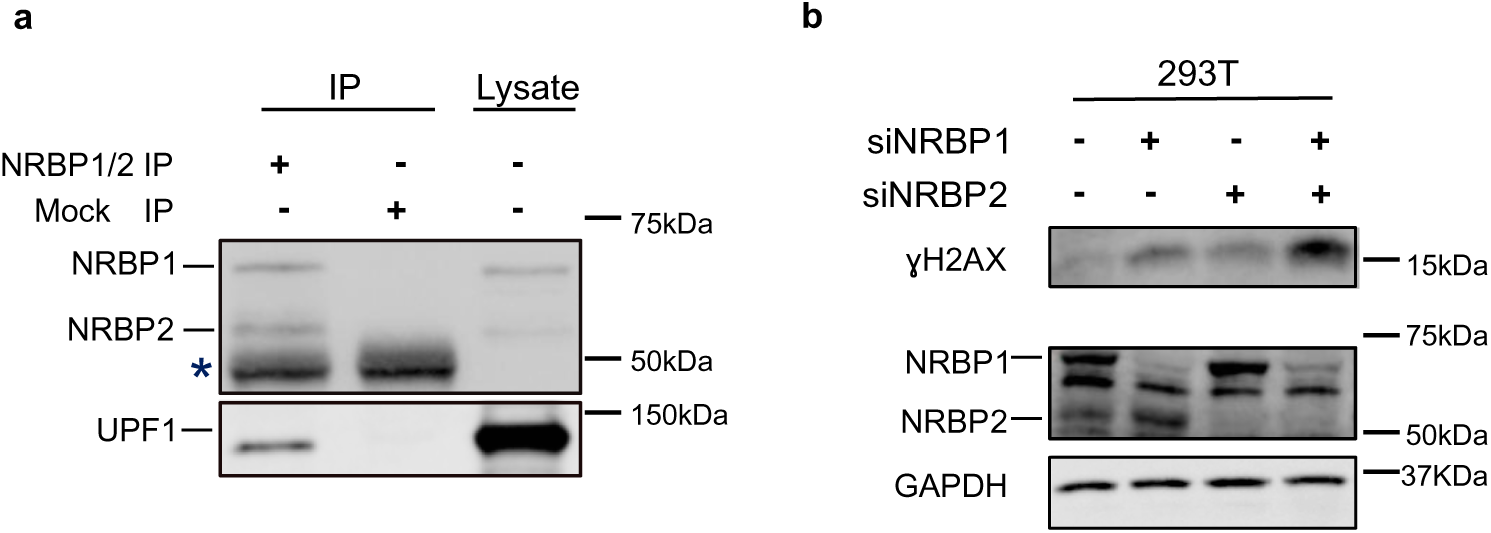
h*p*o*-11* and its human homologs NRBP1 and NRBP2 share conserved functions a. NRBP1/2 interact with UPF1 in MCF-7 cells. Shown is one representative result of three biological replicates. b. siRNA knock-down of either NRBP1 or NRBP2 caused DNA damage response. Shown is a western blot of γ-H2AX in HEK293T cell, N = 3 biological replicates.

## Discussion

In this study, we have discovered an evolutionary conserved function of pseudokinase HPO-11 and its mammalian homologs NRBP1/NRBP2 in ensuring genome stability. Our data also imply that, in the absence of HPO-11, GIN has distinct consequences in the germline and in the soma. DNA damage caused by loss of *hpo-11* results in cell death in the apoptosis competent germline. Efficient removal of germ nuclei carrying DNA damage might explain why *hpo-11* loss of function mutant does not obviously behave like a mutator mutant. In apoptosis deficient somatic cells, however, *hpo-11* knock-down results in elevated mutation rate.

Our work suggests HPO-11 as a negative regulatory of NMD. The factors in NMD were discovered mostly via genetic screens in yeast and *C. elegans* by screening for mutations or RNAi resulting in stabilization of PTC containing mRNAs^40, 47–49^. As those genetic screens only identified positive regulators of NMD, much fewer details are known about how NMD activity is negatively regulated. UPF3A in vertebrates antagonizes UPF3B activity through sequestering UPF2 away from UPF1^50^. In addition, NMD can be inhibited by placing poly(A) binding protein (PABP) in proximity to the termination codon^51^, suggesting that the decreasing distance between the translation termination event and PABP binding antagonizes NMD. Another study in *C. elegans* showed that elevated NMD activity upon reduced insulin signaling contributes to lifespan extension^52^. How insulin signaling mechanistically inhibits NMD, is still unknown. In addition to its evolutionarily conserved role for RNA surveillance, NMD also regulates the levels of certain physiologic transcripts^53^. Given that only a small portion of RNA transcripts bound by SMG-2/UPF1 is indeed down-regulated by NMD, mechanisms contributing to discriminating RNA degradation from RNA-binding are indispensable to avoid unselected, excessive RNA decay. Only 6 out of 32 targets which are regulated by *hpo-11* in a *smg-2* dependent manner, are known NMD substrates (Supplementary Fig. 5b). This result implies that HPO-11 might affect target selection of NMD although it does not harbor any predicted RNA binding motif. As a pseudokinase, HPO-11 might fulfil its regulatory role as an adaptor protein to affect protein interactions and complex associations. Our observation is that interaction between the recombinant HPO-11 and SMG-2 proteins does not apparently depend on RNA (Fig. 4c), suggesting a direct physical binding of HPO-11 to SMG-2. On the other hand, the MS HPO-11 interactor screen failed to identify other components of the NMD complex, like SMG-1 or SMG-3 to SMG-7, as HPO-11 interactors, whereas mutations of *smg-1* to *smg-7* all interfered with HPO-11 function. These results together raise the question whether HPO-11 could act as a sequestering factor to prevent loading of SMG-2 into the NMD complex. In such a scenario, HPO-11 should have a general inhibitory effect on NMD. This is, however, not supported by our transcriptome results. Interestingly, we found endogenous germline HPO-11 formed cytoplasmic foci, which are probably mRNA containing ribonucleoparticles (mRNPs). Therefore, binding of HPO-11 to SMG-2 might rather modulate the recognition of the specific substrates by SMG-2 instead of preventing complex assembly of the canonical NMD machinery. How could HPO-11 specifically select mRNAs for protection? Our MS data indicate that HPO-11 also associates with four proteins of the polyadenylation complex (CPF-1, CPF-2, SUF-1 and CPSF-1) that could affect length of 3’UTR of mRNA by alternative polyadenylation^54^. mRNAs containing long 3′UTRs constitute a major class of physiological targets of NMD without PTC, and sensing the length of 3′UTR by NMD is mediated by UPF1^55^. Therefore, HPO-11 might govern alternative polyadenylation to affect length of 3’UTR which in turn controls substrates specificity of NMD. The precise contribution of HPO-11 to GIN and its interaction with alternative polyadenylation awaits further exploration.

Studies of UPF1, the SMG-2 orthologue, also demonstrated that it associates with S phase progression and genome stability but in a NMD-independent manner^6^. A very recent study has revealed contribution of UPF1 in R-loop formation which in turn stimulates telomere fusion at chromosomal ends^56^. This UPF1 function is, however, independent of NMD. In our study, the reduced number of chromosomes in the oocytes of *hpo-11(lf)* animals may also be caused by chromosome fusion, but in a both SMG-2 and NMD dependent manner, suggesting a similar biological role of SMG-2/UPF1 that depends on different molecular mechanisms.

Both NRBP1 and NRBP2 have been recently reported to affect tumorigenesis and progression^12–14, 17, 57–59^. In these studies, NRBP1 and NRBP2 can act either as tumor promoter or suppressor, suggesting a tumor-type dependent role in affecting cell survival and proliferation. While molecular functions of NRBP2 are poorly known, NRBP1 is well characterized as a substrate recognition factor within the Elongin BC ubiquitin ligase complex to regulate proteasome mediated degradation of specific proteins^17^. In addition, NRBP1 is reported to interact with JAB1 to inhibit AP1 which activates transcription of several DNA repair genes in response to genotoxic damage^60, 61^. Our studies demonstrate that the *C. elegans* HPO-11 as well as its mammalian homolog NRBP1/2 have a crucial role in genome maintenance that is probably independent of the Elongin BC ubiquitin ligase complex. HPO-11 rather protects genome integrity by adjusting proper NMD activity to prevent the degradation of specific transcripts. Our findings not only propose a potential interplay between NMD and DNA damage, but also identify HPO-11 as a negative modulator of NMD to prevent GIN. In eukaryotes, maintenance of genome integrity is essential for ensuring genome function and for precluding mutations that might lead to tumorigenesis. On the other hand, tumor cells frequently suffer from increased replication stress resulting from elevated levels of DNA damage. Consequently, malignant cells become even more dependent on DNA repair mechanisms to survive and proliferate. Conventional tumor treatment such as radiation therapy and certain forms of chemotherapy induce DNA damage to provoke cell death. Because HPO-11 functions in genome stability maintenance is evolutionary conserved, deciphering its mechanisms in *C. elegans* will shed light on the functions of NRBP1/2 in tumor biology and suggest them as potential targets in cancer therapy.

## Materials and Methods

### Strains

The *C. elegans* N2 (Bristol) strain was used as wild type in all experiments in this study. BR7508 *hpo-11(by178)*, BR7844 *hpo-11(by208[YPET::LoxP::3xFlag::hpo-11])*, BR8845 *hpo-11(by233)* (a *hpo-11* rescue strain that repairs the mutation in *by178* via CRISPR-Cas9 mediated genome editing). IJ445 *smg-2(qd101)*, BR8580 *smg-2(qd101) hpo-11(by178)*, MD701 *bcls39[Plim-7::ced-1::gfp;lin-15(+)]*, BR8837 *hpo-11(by178);bcls39[Plim-7::ced-1::gfp; lin-15(+)]*, NL2250 *pkIs1604[hsp-16.2P::ATG(A)17GFP::LacZ + rol-6(su1006)]*, BR8681 *hpo-11(by178);pkIs1604[hsp-16.2P::ATG(A)17GFP::LacZ + rol-6(su1006)]*, *Is[sec-23p::gfp::lacZ(PTC);unc-76],* BR8584 *hpo-11(by178);Is[sec-23p::gfp::lacZ(PTC);unc-76],* BR8430 *Ex1762[hpo-11P::YPET::LoxP::3xFlag;rol-6(su1006)]* (as IP control in the MS analysis), BR5082 *shc-1(ok198);Is[daf-16::GFP]*, BR7037 *shc-1(ok198) hpo-11(by178);Is[daf-16::GFP].* BR6826 *byIs212[hpo-11P::hpo-11(wt)::EGFP;rol-6(su1006)]* BR8951 *byEx2665[hpo-11P::hpo-11(by178)::EGFP,myo-2P::mCherry]*.

### Plasmids and cloning

To express recombinant proteins, *smg-2* cDNA was inserted into pcDNA6/V5-6xHis vector at *NheI/EcoRI* sites to obtain pBY4140 (*pcDNA6A::smg-2::V5::6xHis*), *hpo-11* cNDA was inserted into pEGFP-C1 at *SacI/KpnI* sites to obtain pBY4139 (*pEGFP-C1::hpo-11*). pBY4044 *hpo-11P:: YPET::LoxP::3xFlag* contains the 3676 bp *hpo-11* promoter region upstream of translational start region and a 1038bp fragment carrying *YPET::LoxP::3xFlag* sequence that was amplified from BR7844 *hpo-11(by208[YPET::LoxP::3xFlag::hpo-11].* The two fragments were inserted into pEGFP-C1 with *Eco47III/AgeI* and *AgeI/KpnI* sites, respectively. To obtain pBY3725 *hpo-11P::hpo-11(wt)::EGFP*, 3672 bp *hpo-11* promoter region and 6955 bp of *hpo-11* coding region were amplified from wild type DNA and inserted into pEGFP-N1 vector with *Eco47III/SacI* and with *SacI/AgeI* sites, respectively. To obtain pBY4202 *hpo-11P::hpo-11(by178)::EGFP*, the 1701 bp fragment of pBY3725 were removed with SacI/BamHI digestion and replaced with the same DNA fragment amplified from *hpo-11(by178)* genomic DNA.

### Quantification of embryonic viability

L4 hermaphrodites were picked to individual plates and transferred to new plates for egg laying daily until day 3 adulthood. Total number of laid eggs and hatched larvae were counted. The percentage of viable progeny was calculated by dividing the number of hatched progenies through the number of total eggs.

### Quantification of sex ratio

L4 hermaphrodites were picked to individual plates and transferred daily to new plates for egg laying for 3 days. Male and hermaphrodite progeny were counted when animals reached L4 stage. The percentage of male progeny was calculated by dividing the number of the male progeny by the total number of progeny.

### RAD-51 and R-loops antibody staining

The RAD-51 antibody staining was performed with extruded gonads of day one adult worms. Animals were picked to 50 µl M9 buffer containing 0.25 mM levamisole and gonads were extruded by cutting head and tail with surgical blades. Dissected gonads were fixed with 1% formaldehyde for 10 minutes at room temperature in a humid chamber followed by 1 h incubation in PBST-A (1% BSA, 1xPBS, 0.1% Tween 20 and 0.05% NaN3). The samples were washed with 500 µl PBST-A and suspended in 5 µl PBST-A before antibody staining. 10 µl of 1:50 diluted mouse anti-RAD-51 antibody (diluted 1:50 in PBST-A) were added to the worms for incubation overnight at 4 °C. After 3 times subsequent washing for 15 minutes each with PBST-B (0.1% BSA, 1xPBS, 0.1% Tween20 and 0.05% NaN3), the samples were incubated with secondary antibody Cy3 anti-mouse (Jackson Immuno research, 715-166-150,1:200 dilution in PBSTA) for 2 hours at room temperature. The gonads were washed 3 times and then stained with 1 μg/mL 4’,6-diamidino-2-phenylindole (DAPI) before mounting with antifade solution (Invitrogen, P36980) on poly-L-lysine coated glass slides (Sigma Merck, P0425-72EA). The edges of the slides were sealed with nail polish. Images were taken using a Leiss LSM880 confocal microscopy with 40x/1.4 Oil objective. Confocal images shown are maximum intensity projections of 0.5–0.8 µm Z-stacks of the entire gonad. For display, contrast and brightness were adjusted in individual color channels in ZEN (black edition). At least five gonads were counted per genotype for quantification of RAD-51 foci.

R loops were stained with an S9.6 antibody that recognizes DNA-RNA hybrid (MABE1095, Sigma-Aldrich) by using the same staining and confocal images procedure as RAD-51 antibody staining. Quantification of R loops foci was performed by dividing the number of total stained S9.6 foci by total number of germ nuclei in the early pachytene region. At least five gonads were counted per genotype.

### Fluorescence microscopy

*hpo-11(by208[YPET::LoxP::3xFlag::hpo-11])* worms were immobilized with 0.25 mM levamisole on 2% agarose pads and imaged using a LSM-U-NLO confocal microscope (Carl Zeiss) with a 40x objective. Images were adjusted as necessary in ZEN black using cropping, brightness and contrast tools. Subcellular localization of YPET::LoxP::3xFlag::HPO-11 in response to CPT treatment was imaged by using an Imager.Z1 microscope (Carl Zeiss). For visualizing chromosomes in the oocytes, dissected gonads were fixed in methanol and re-suspended in PBST containing 0.1% Tween 20 and 2 μg/ml DAPI before microscopy. Z-stack images of the gonads were collected with the Imager.Z1 compound microscope for counting numbers of the chromosomes in the most proximal oocyte.

### X-gal staining

Day 1 adult worms were washed in distilled water and frozen in liquid nitrogen. After dried in a vacuum jar for 45 minutes, worms were soaked in cold acetone at -20°C for 3 minutes. Liquid was removed. For ß-galactosidase detection, worms were incubated in 1 ml freshly made X-gal staining solution (620 μl ddH_2_O, 200 μl 1 M pH 7.5 Na-phosphate buffer, 1 μl 1 M MgCl_2_, 4 μl 1% SDS, 50 μl 100 mM Potassium Ferricyanide, 50 μl 100 mM Potassium Ferrocyanide, 8 μl 5% X-Gal in dimethyl formamide)

### Anti-FLAG co-immunopreciptiation and in-gel digestion

*hpo-11(by208[YPET::LoxP::3xFlag::hpo-11])* and *Ex1762[hpo-11P::YPET::LoxP::3xFlag;rol-6(su1006)]* (as mock IP) worm samples were collected from mixed stages, washed three times with 1x ice cold PBS and immediately frozen in liquid nitrogen. Worms were lysed in 800 µl Triton-X-100 lysis buffer (150 mM NaCl, 50 mM Tris pH 8.0, 1% Triton-X-100 + Protease Inhibitor+ phosphatase inhibitor) with SilentCrusher S (90 s, 12,600 x g). Lysates were cleared by centrifuging at 10,000 g for 15 min at 4 °C. 100 μl supernatant was taken for concentration measurement as well as analyzed for input. Next, supernatant containing 300 μg protein was pre-cleaned with 100 μl protein G dynabeads (Thermo Fisher Scientific Biosciences, 10009D) for 30 min at 4°C on the overhang shaker. 50 μl Anti-FLAG beads (Sigma, M8823-1ML) were washed three-times with lysis buffer before incubation with 50 μl protein G dynabeads. Pre-cleaned supernatant was eluted with 100 µl 1x lysis buffer.

Elutes were loaded onto NuPAGE® Novex® 4–12% Bis-Tris gels (Invitrogen, Life Technologies). After gel electrophoresis, proteins were stained with Coomassie Brilliant Blue G-250. Ingel protein digestion using trypsin (Promega, Mannheim, Germany) was performed essentially as described before^62^. Proteins were destained using 50% ethanol in 10 mM ammonium bicarbonate. For reduction of disulfide bonds and subsequent alkylation, 5 mM tris(2-carboxyethyl) phosphine (10 min at 60 °C) and 100 mM 2-chloroacetamide (15 min at 37 °C) were used, respectively. Proteins were digested with trypsin in 10 mM ammonium bicarbonate at 37°C overnight and peptides were extracted using one volume of 0.1% trifluoroacetic acid in water mixed with one volume of 100% acetonitrile. The peptides were dried down and taken up in 15µl 0.1% trifluoroacetic acid in water.

### High-Performance Liquid Chromatography and Mass Spectrometry (HPLC-MS)

HPLC-MS analysis was performed either on an Ultimate^TM^ 3000 RSLCnano system coupled to an LTQ Orbitrap XL (two replicates) or a Velos Orbitrap Elite instrument (one replicate). All instruments are from Thermo Fisher Scientific, Bremen, Germany. On both HPLCs a binary solvent system was used with solvent A consisting of 0.1% formic acid and 4% DMSO and solvent B consisting of 48% methanol, 30% acetonitril, 0.1% formic acid and 4% DMSO. The HPLC coupled to the LTQ Orbitrap XL was equipped with two PepMapTM C18 μ-precolumns (ID: 0.3 mm × 5 mm, 5 µm, 300 Å Thermo Fisher Scientific) and an Acclaim^TM^ PepMap^TM^ analytical column (ID: 75 μm × 250 mm, 3 μm, 100 Å, Thermo Fisher Scientific). Samples were washed and concentrated for 5 min with 0.1% trifluoroacetic acid on the pre-column. A flow rate of 0.250 µl/min was applied to the analytical column and the following gradient was used: 1% B to 30% B in 34 min, to 45% B in 12 min, to 70% B in 14 min, to 99% B in 5 min, 99% B for 5 min and decreased to 1% B in 1 min. The column was re-equilibrated for 19 min with 1% B. The HPLC coupled to the Velos Orbitrap Elite instrument was equipped with two nanoEase™ M/Z Symmetry C18 trap columns (100Å pore size, 5µm particle size, 20 mm length, 180 µm inner diameter) and a nanoEase™ M/Z HSS C18 T3 analytical column (250 mm length, 75 µM inner diameter, 1.8 µM particle size, 100 Å pore size), all from Waters Corporation, Milford, MA. The trap columns were operated at a flow rate of 10 µl/min and the analytical column at 300 nl/min. Peptide samples were pre-concentrated on the trap column using 0.1% trifluoroacetic acid for 5 min before switching the column in line with the analytical column. Peptides were separated using a liner gradient, with an increase of solvent B from 3% to 55% in 65 min, followed by an increase to 80% B in 5 min. The column was washed for another 5 min with 80% B before returning to 3% B in 4 min and re-equilibration for 21 min.

The MS instruments were operated with the following parameters: 1.5 kV spray voltage, 200°C capillary temperature. Orbitrap mass range on both instruments’ m/z 370 to 1,700. For the LTQ Orbitrap XL the resolution at *m*/*z* 400 was 60,000, automatic gain control 5×10^5^ ions, max. fill time, 500 ms. For the Velos Orbitrap Elite, the resolution at m/z 400 was 120,000, automatic gain control 1×10^6^ and the maximum ion time 200 ms. A TOP5 (LTQ Orbitrap XL with automatic gain control 10,000 ions, max. fill time 100 ms) or a TOP 25 (Velos Orbitrap Elite with automatic gain control 5,000, max. fill time 150 ms) method was applied for collision-induced dissociation of multiply charged peptide ions. On both instruments, the normalized collision energy was 35% and the activation Q 0.250. Dynamic exclusion was set to 45 s.

### MS Data analysis

Raw files were searched with MaxQuant version 1.6.10.43^63, 64^ against the *C. elegans* UniProt reference proteome (ID: UP000001940; Taxonomy: 6239; 26984 protein entries; 5.11.2019), complemented with the YFP-FLAG-tag amino acid sequence.

Over all, default settings were used in MaxQuant. Trypsin/P was used as proteolytic enzyme and up to two missed cleavages were allowed. 1% false discovery rate was applied on both peptide and protein lists. Methionine oxidation and N-terminal acetylation were set as variable and carbamidomethylation as fixed modifications. Minimum number of unique peptides was set to 1.Label-free quantification^65^ was enabled, with a minimum ratio count of two and the option “require MS/MS for label-free quantification (LFQ) comparisons” enabled.

For data analysis, the proteingroups.txt file of MaxQuant was used and loaded into Perseus 1.5.5.3^66^. Entries for reverse and contaminant hits as well as proteins only identified by site were removed from the analysis. LFQ intensities were log_2_-transformed. Only protein groups with LFQ intensities either in all three replicates of the FLAG-HPO11 IP or in all three replicates of the corresponding control IPs were considered for further analysis. Missing values were imputed from normal distribution using the following settings in Perseus: width 0.5 and down shift 1.8. A one-sided two sample t-test was performed. Proteins with a p-value below 0.05 and a student’s t-test difference of at least 2 were considered as significantly enriched candidates.

The MS data have been deposited to the ProteomeXchange Consortium (http://proteomecentral.proteomexchange.org) via the PRIDE partner repository^67^ with the dataset identifier PXD027876.

### Gene Ontology Analysis

Gene ontology analysis was performed on the interaction partners of HPO-11, using DAVID v.6.8 (https://david.ncifcrf.gov) for classifying based on biological process and molecular function.

### RNA isolation and sequencing

Synchronized day one adult animals of N2, *hpo-11(by178), smg-2(qd101)* and *smg-2(qd101) hpo-11(by178)* grown up at 20 °C were washed off from NGM plates using M9 solution and subjected to total RNA extraction using the RNeasy Mini Kit from Qiagen. Purification of poly-A containing RNA molecules, RNA fragmentation, strand-specific random primed cDNA library preparation and single-read sequencing (50 bp) on an Illumina HiSeq 4000 were carried out by Eurofins Genomics. Three biological replicates of each genotype were sequenced.

### RNA-sequencing data analysis and graphic generation

FastQ files were uploaded to Galaxy/Europe (https://usegalaxy.eu/). FastQC was used for the quality control of raw sequencing data^68^. Trim galore was applied to trim the low quality reads of 3’ prime and 5’ prime ends. Pre-treated reads were mapped with RNA star^69^(Galaxy Version 2.7.2b) to *C. elegans* reference genome (WBcel235.96) from Ensembl. Aligned rate was >93% with >30 million uniquely aligned reads. The expression levels of transcripts were measured with feature counts^70^ (Galaxy Version 2.0.1). DESeq2 (1.30.1) R package was used for estimating the differential expression level of transcripts based on experiment design^71^.

### Quantitative Real-Time PCR (qRT-PCR)

Total RNA was prepared from synchronized day one adult animals raised at 20°C by using RNeasy® Mini Kit (Qiagen, Venlo, The Netherlands). Oligo-dT primer was used to synthesize cDNA by using Transcription First Strand cDNA synthesis kit (Roche) according to product user guide. qRT-PCR was performed in the Light Cycler 96 System (Roche) using Luna Universal qPCR Master Mix (NEB). Primers for *rpl-12(PTC)* and *rpl-12(wt)* were taken from a previous study^52^. The other primers for qRT-PCR were designed in Primer-Blast (https://www.ncbi.nlm.nih.gov/tools/primer-blast/) or manually selected based on the following parameters: (1) the amplicon is between 70 and 200 bp long; (2) the melting temperature is 60°C; and (3) primers must be across an exon-exon junction. Data were processed with relative quantification 2^-ΔΔCt^ method^72^ and normalized to *act-4*. All qRT-PCR primers used in this study are shown in Supplementary Table 1.

### Measurement of embryonic survival upon camptothecin (CPT) treatment

Animals in L4 stage were transferred to NGM plates containing 5 μM CPT for 24 h and then transferred to CPT-free plates with for another 24 h to allow egg laying. 24 h after removal of the adult animals, the number of hatched and unhatched eggs was counted. The percentage of embryonic survival was calculated by dividing the number of hatched eggs by the total number of eggs laid.

### Co-immunoprecipitation (Co-IP)

For Co-IP of recombinant EGFP::HPO-11 and V5::SMG-2, HEK 293T cell line was cultured in Dulbecco’s modified Eagle’s medium (DMEM), supplemented with 10% fetal bovine serum and 1.5 % L-glutamine Cells were plated one day before transfection to ∼ 60% confluency and transfected using jetPEI (Ployplus) according to the manufacturer’s protocol. 48 h after transfection, cells were washed three times with cold 1x PBS collected with Echelon lysis buffer (20 mM Tris-HCL, pH 7.4, 137 mM NaCl, 1mM CaCl_2_, 1mM MgCl_2_, 1% (v/v) NP-40, 1 mM Na_3_VO_4_, freshly added 1mM PMSF) followed by centrifugation at 10,000 *g* for 15 min at 4 °C. The supernatant was incubated with protein A magnetic beads on a rotator for 30 minutes at 4°C. 300 μg total protein was used for immunoprecipitation with 0.25 μg anti-GFP antibody on rotator overnight at 4°C. Protein A magnetic beads were added into the antibody-antigen mixture followed by incubation for 2 h at 4 °C. Beads were then collected with a magnetic stand and washed three times in wash buffer. Protein complex was eluted from beads with 1x Laemmli buffer. Input and eluted IP samples were analyzed by western blot.

Immunoprecipitation (IP) of endogenous NRBP1 and NRBP2 was performed in MCF7 cells. Cells were cultured in RPMI 1640 (Fisher Scientific) with 10% FBS (Sigma-Aldrich) and 2 mM glutamine and subdivided into four times 15 cm dishes for IP of target protein and mock IP, respectively. Dishes were washed twice in cold PBS and PBS was removed. Then cells were lysated with 1.5 ml 1x Sabatini lysis buffer (Phosphatase Inhibitor Cocktail 2 (P5726, Sigma), Phosphatase Inhibitor Cocktail 3 (P0044, sigma) and Protease Inhibitor Cocktail (11697498001, sigma)). Scraped off cells were transferred to 2 ml Eppendorf tubes for overhead rotation for 30 minutes at 4°C. Supernatant was incubated with pre-cleaned protein G dynabeads (Thermo Fisher Scientific Biosciences, 10009D) for overhead rotation for 30 minutes at 4°C. Antibodies for IP (NRBP1/2 rabbit, 21549-1-AP –ptg) and mock IP (IgG rabbit, Santa Cruz Biotech, sc-2027) were then added into the supernatant. The IP was performed for 1 h at 4°C on an overhead rotator before incubation with pre-cleaned protein G dynabeads. Proteins were eluted from beads with 1x Laemmli buffer. Anti-UPF1 antibody (1:1000 dilution, D15G6-CST from Cell Signaling Technology) was used to detect in the IP group and mock group by western blot.

### Detection of γH2AX upon siRNA knock-down of NRBP1 and NRBP2

All ON-TARGET plus siRNA were purchased from Dharmacon (Horizon Discovery Ltd). HEK293T cells were transfected using Lipofectamine™ RNAiMAX Transfection Reagent (Thermo Fisher Scientific Biosciences). The siRNAs were diluted in 500 µl Opti-MEM®I Medium without serum in 6-well tissue culture plates. 3 µl Lipofectamine RNAi MAX was added to each diluted siRNA. After incubation for 20 minutes at room temperature, the siRNA-Lipofectamine mixture was added into each well with the concentration of 250,000 cells/ml. Transfected cells were incubated for 72 h before subjecting to western blot analysis. First, knockdown efficiency of NRBP1/2 was determined using anti-NRBP1/2 antibody. The phosphorylated H2AX was detected using anti-phospho-histone H2AX (Ser139) (20E3) (1: 1000 dilution, 9718T from Cell Signaling Technology).

### Antibodies

Anti-Flag (F3165, Sigma-Aldrich), anti-V5-Tag antibody (ab27671, Abcam), anti-GFP antibody (ab290, Abcam), Anti-DNA-RNA hybrid antibody S9.6 (MABE1095, Sigma-Aldrich), anti-NRBP1/2 antibody (21549-1-AP, Proteintech Europe), Phospho-Histone γH2AX antibody (Ser139) (9718, Cell Signaling Technology) and GAPDH (ab8245, Abcam). Anti-RAD-51 antibody is kindly provided by Verena Jantsch, University of Vienna.

### Data availability

The mass spectrometry proteomics data have been deposited to the ProteomeXchange Consortium via the PRIDE^67^ partner repository with the dataset identifier PXD027876, RNA-seq data were submitted to BioProject with ID PRJNA728052. The remaining data for analyzed RNA-seq data are summarized in Supplementary Data 2. Correspondence and requests for materials should be addressed to R.B. (baumeister@celegans) or W.Q.

## Acknowledgements

We thank Dennis Kim and Javier F. Cáceres, Seung-Jae V. Lee and the *Caenorhabditis Genetics Center* providing some of the *C. elegans* strains used in this study. We thank Ruth Jähne for generating six sgRNAs for *hpo-11(by233)* strains. We also thank the staffs of the Life Imaging Center (LIC) of the Albert-Ludwigs-University Freiburg for their microscopy resources (DFG INST 39/1012-1 FUGG). Specially thanks Verena Jantsch (University of Vienna) for the anti-RAD51 antibody. We are particularly grateful to Bettina Warscheid for her careful reading of the manuscript and her valuable suggestions on the MS experiments and their interpretation. This work was funded by grants from the German Research Foundation (DFG) Project ID 403222702/SFB 1381 to R. B. and B. W., SFB850 to RB, Germany’s Excellence strategy (CIBSS-EXC-2189 - Project ID 8 390939984), the German Excellence Initiative (BIOSS - EXC294) to R. B. and B. W. as well as China Scholarship Council to Q. Z.

## Author Contributions

Q.Z., R.B., and W.Q. designed the experiments and analyzed data. Q.Z., E.D.v.G., W.Y., J.S., E.S. designed, B.S. and L.T. performed the *hpo-11* CRISPR knock-in and/or rescue experiments. B. W and J.S. supervised and performed LC-MS analysis, respectively. Q.Z. analyzed RNA-Seq data. Q.Z., W.Q. and R.B. wrote the manuscript.

## Figures and legends

**Supplementary Fig. 1:**
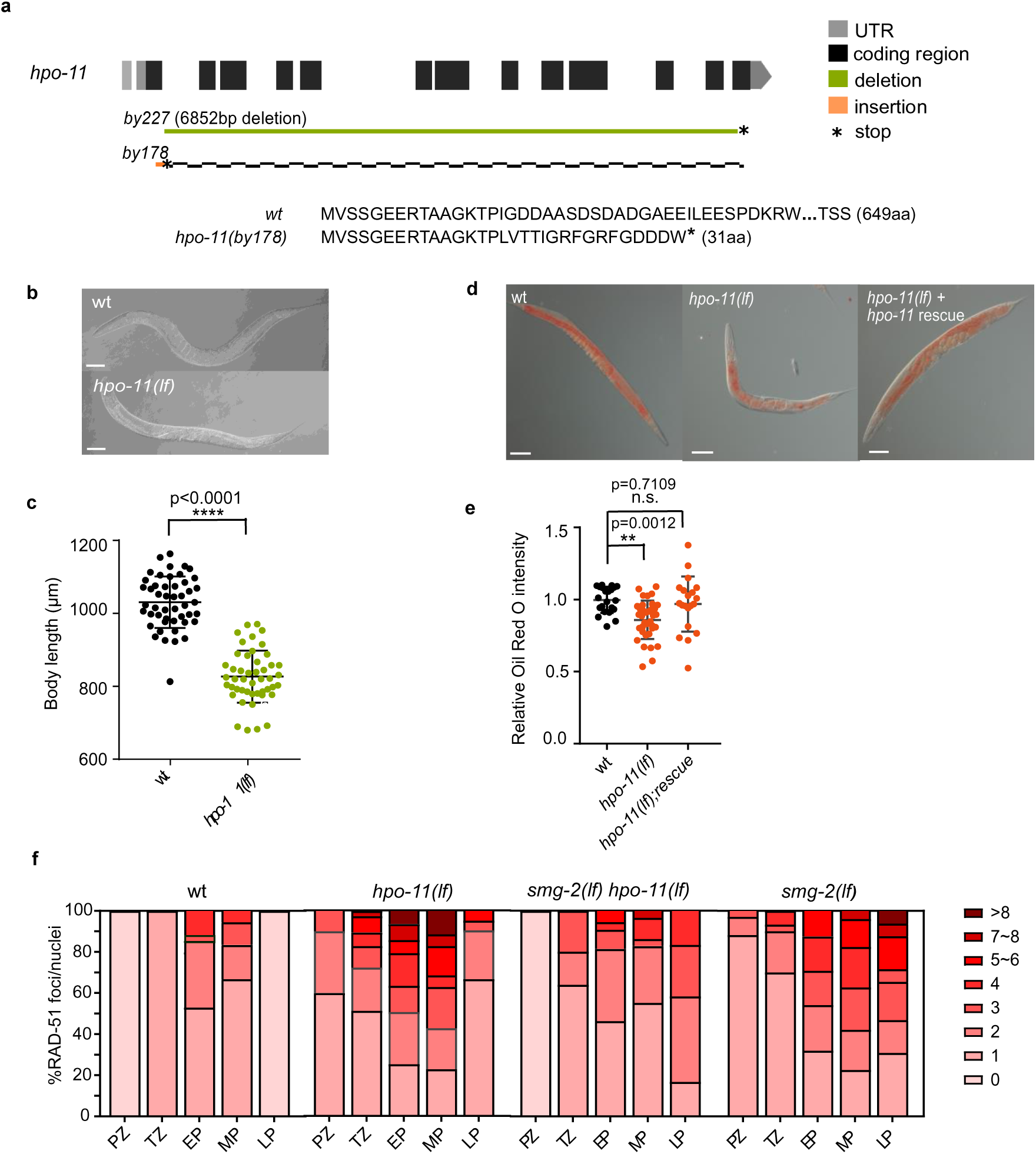
*hpo-11* loss of function mutants display pleiotropic phenotypes. a. Schematic diagram illustrating the *hpo-11(by178)* and *hpo-11(by227)* alleles. A 45 nt insertion in *by178* leads to a premature stop codon in the second exon of *hpo-11* and a frameshift of the downstream sequence. *hpo-11(by227)* contains a 6852 nt deletion which eliminates almost the complete coding region. b. Representative DIC images showing smaller body size of *hpo-11(by178)* animals. Scale bar 50 µm. c. Quantification of the body length of wild type and *hpo-11(by178)* day one adult animals. Wild type n= 48, *hpo-11(by178)* n= 47. Statistical test with two-tailed t test. d. Representative DIC images showing Oil-red-O lipid staining of wild type, *hpo-11(by178)*, and *hpo-11(by233)* rescue day one adult animals. Scale bar 50 µm. e. Quantification of lipid content via Oil-red-O lipid staining in wild type, *hpo-11(by178)*, and *hpo-11(by233)* rescue day one adult animals. Wild type n=21, *hpo-11(by178)* n=34, *hpo-11(by233)* n=20. Shown are pooled data of two biological repeats of the experiments. Statistical test with one-way ANOVA. f. Quantification of RAD-51 foci in the gonad of wild type, *hpo-11(by178)*, *smg-2(qd101)*, *smg-2(qd101) hpo-11(by178)* day one adult animals. At least five gonads were analyzed for each genotype. This figure is related to the main Fig. 1. Supplementary Fig. 1f is also related to the main Fig. 3.

**Supplementary Fig. 2:**
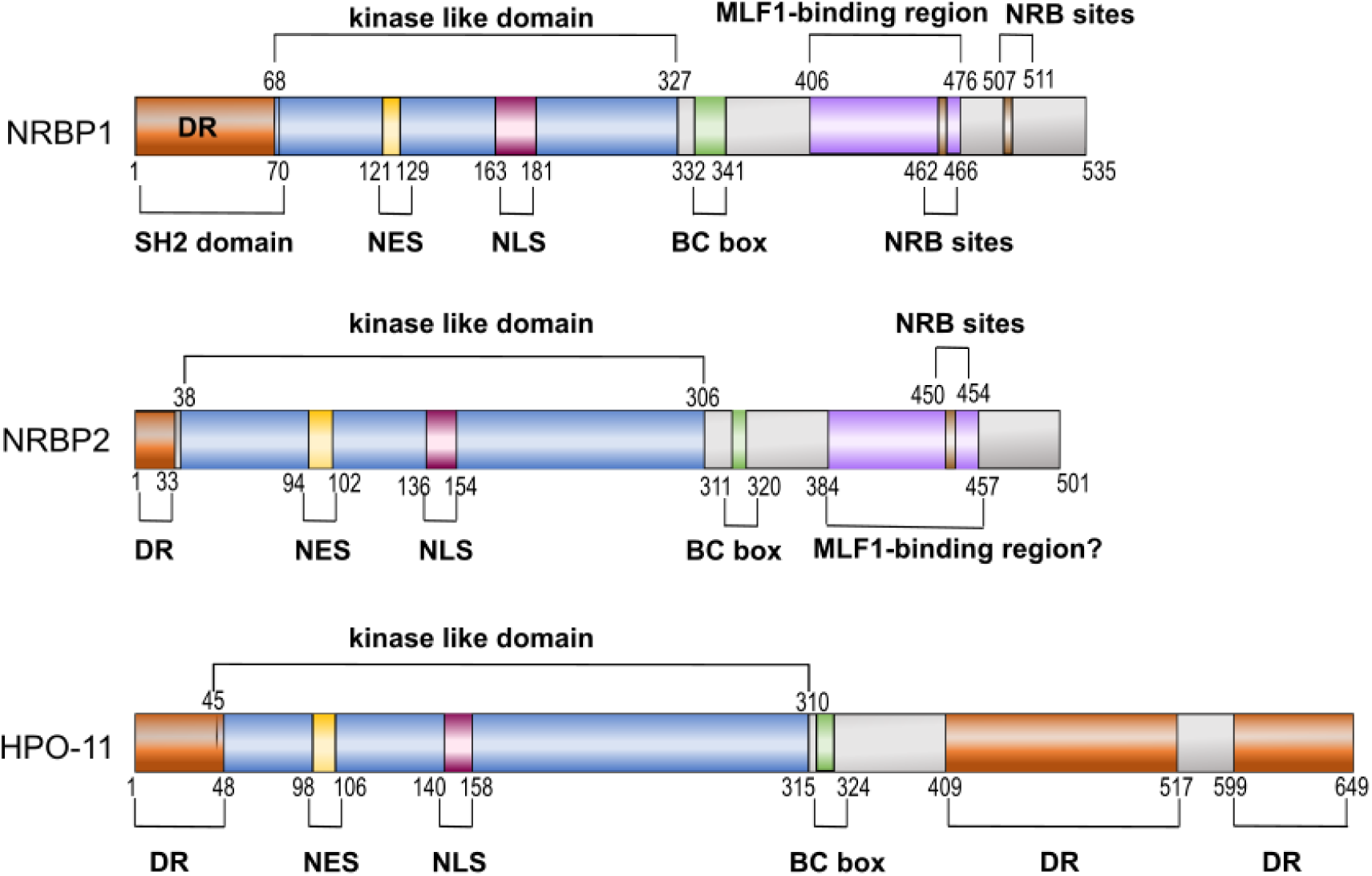
*C. elegans* HPO-11, human NRBP1 and NRBP2 are highly conserved pseudokinases. Shown are the structure domains of the respective proteins. NES: nuclear export signal, NLS: nuclear localization signal, NRB sites: nuclear receptor binding sites, BC box: elongin B/C binding sequence, DR: disordered region. This figure is related to the main Fig. 2.

**Supplementary Fig. 3:**
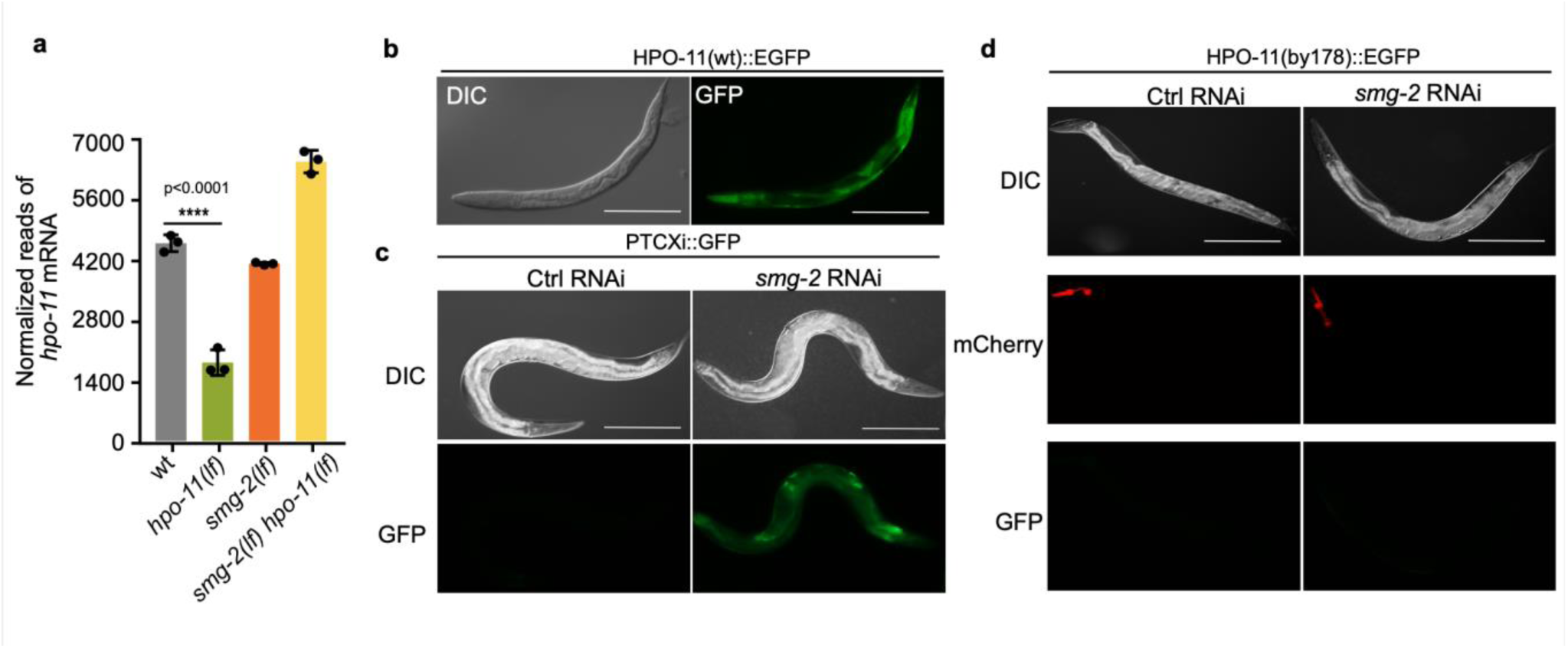
Stabilization of *hpo-11(by178)* mRNA by inactivating *smg-2* does not lead to translation of a functional HPO-11(by178) protein. a. RNA-seq results of *hpo-11* mRNA level in different strains. *hpo-11(by178)* has significantly reduced PTC containing *hpo-11(by178)* mRNA level that is increased by additional inactivating *smg-2*. b. Expression of *hpo-11(wt)::GFP* reporter in day one adult animals. Scale bar 200 µm. c. *smg-2* RNAi activates expression of a PTC containing reporter PTCxi::GFP that does not have a frameshift of GFP in day one adult animals. Scale bar 200 µm. d. *smg-2* RNAi does not activate expression of *hpo-11(by178)::EGFP* expression in day one adult animals. A myo-2P::mCherry reporter was used as co-injection marker. Scale bar 200 µm. This figure is related to the main Fig. 3.

**Supplementary Fig. S4:**
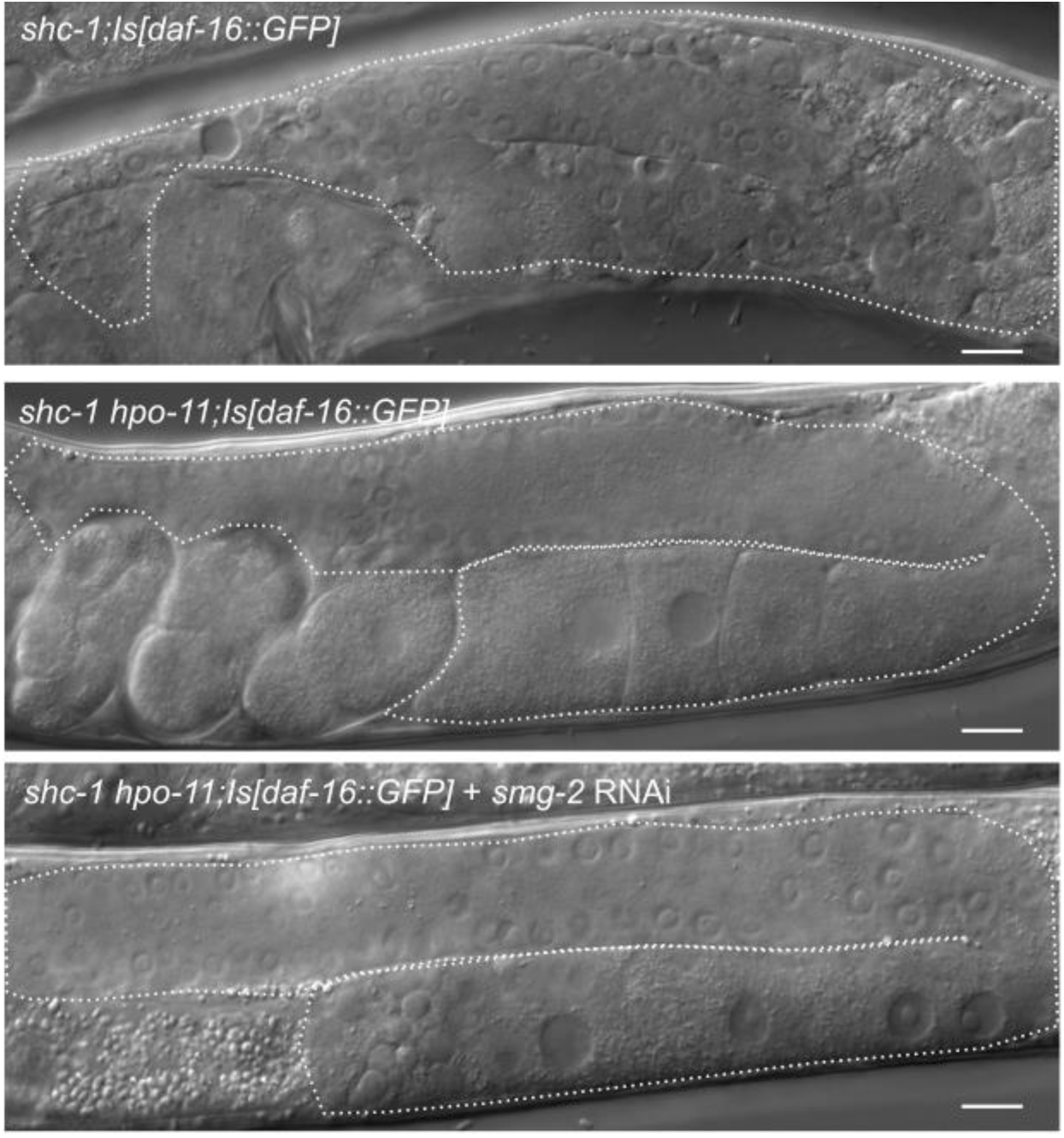
HPO-11 promotes germline tumor formation independent of SMG-2. Shown are DIC images of *shc-1(ok198);Is[daf-16::GFP]*, *shc-1(ok198) hpo-11(by178);Is[daf-16::GFP]* day one adult animals fed with L4440 or *smg-2* RNAi. *shc-1(ok198);Is[daf-16::GFP]* animals display germline hyperproliferation and degradation of the gonadal basement membrane that leads to fill-up of the body cavity with germ nuclei. *hpo-11* knock-down suppresses the germline tumor formation and additional *smg-2* RNAi knock-down does not result in reoccurrence of the germline tumor phenotype. Scale bar 10 µm.

**Supplementary Fig. S5.**
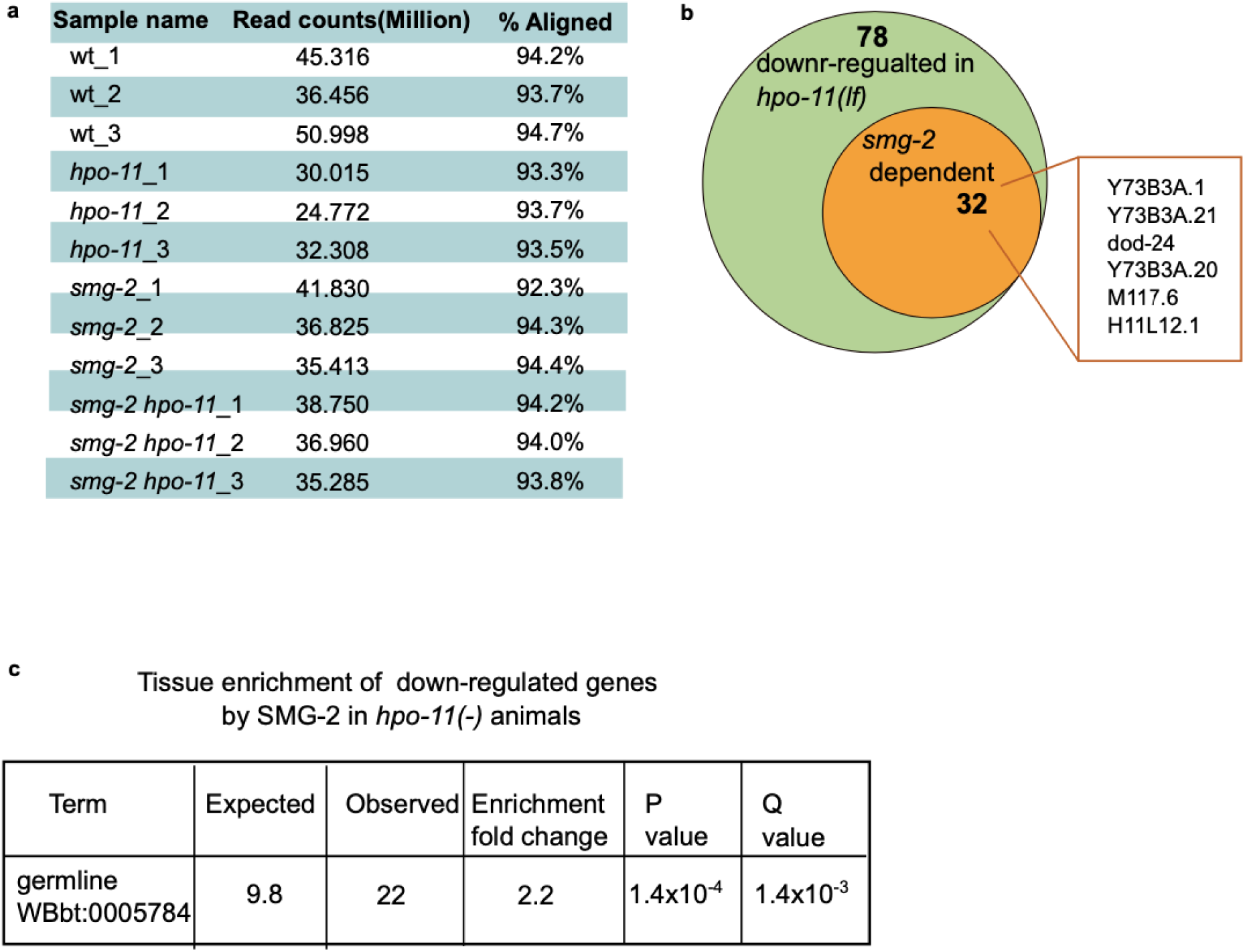
RNA-seq results. a. Summary of sequenced as well as aligned read counts of the all samples. b. HPO-11 negatively regulates, *smg-2* dependent and reported NMD target genes. c. Tissue enrichment analysis of targets controlled by *hpo-11* in *smg-2* dependent manner. This figure is related to the main Fig. 5.

**Supplementary Fig. S6.**
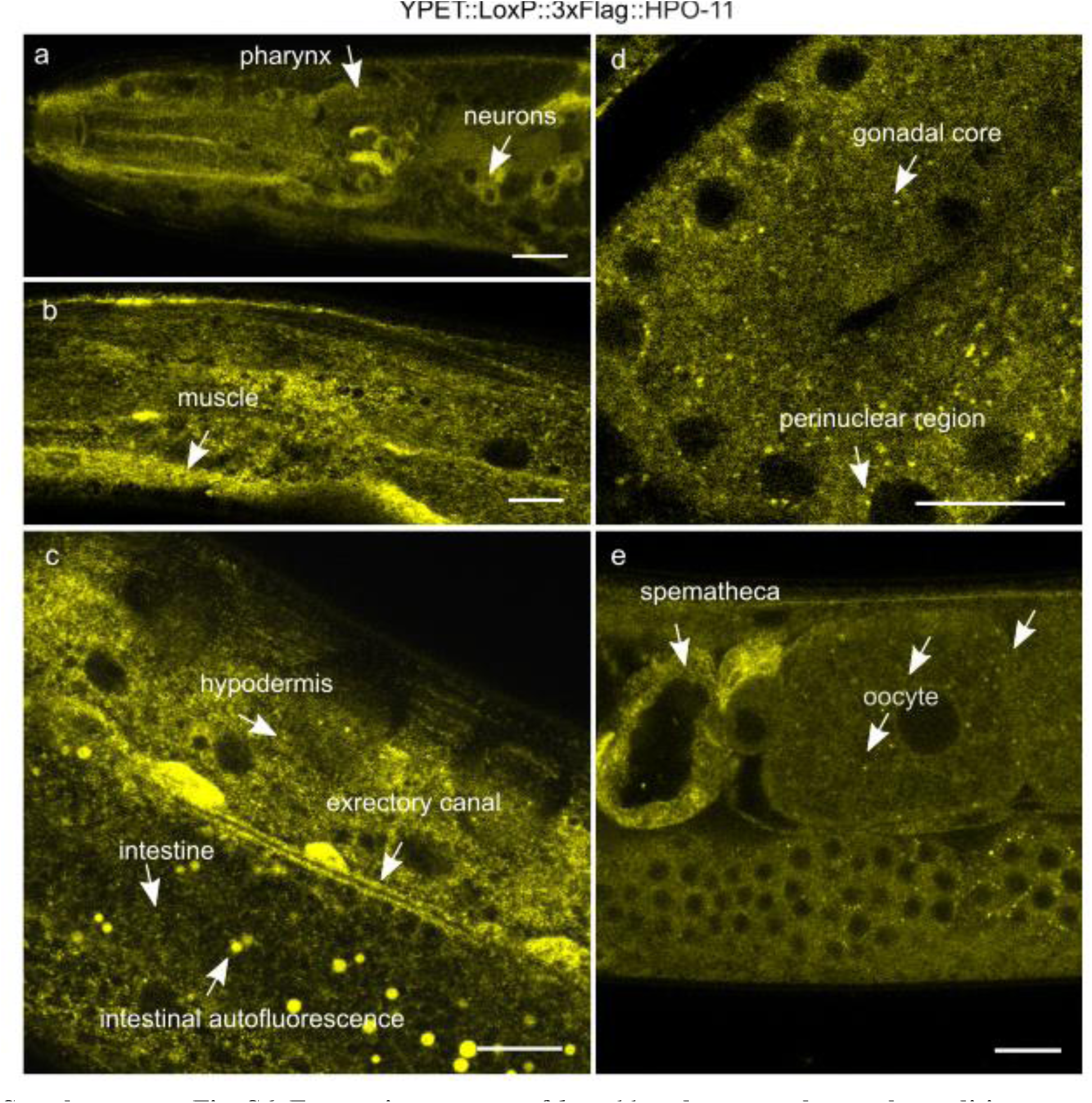
Expression pattern of *hpo-11* under normal growth condition. Shown are fluorescence micrographs of the CRISPR-knock in strain *hpo-11(by208[YPET::LoxP::xFlag::hpo-11])*. YPET::LgoxP::3xFlag::HPO-11 protein is wildly expressed both in the soma and the germline. Scale bar 10 µm. This figure is related to the main Figure 6.

